# G-quadruplexes regulate chromatin accessibility and gene expression in Bloom Syndrome

**DOI:** 10.64898/2026.03.03.709275

**Authors:** Dingwen Su, Veronika Altmannova, Volker Soltys, Moritz Peters, Christopher M Cunniff, John R Weir, Yingguang Frank Chan

## Abstract

Bloom Syndrome (BS) is a recessive genetic disorder characterized by hyper-recombination and genome instability. It is caused by mutations in *BLM,* which encodes a conserved RecQ helicase that unwinds various aberrant DNA structures. One such structure is DNA G-quadruplexes (G4s), which have versatile regulatory potential in chromatin organization and gene expression. However, whether G4 profiles are altered in BS and how G4s contribute to disease-associated molecular changes remain unclear. Here, we profiled chromatin accessibility and gene expression using ATAC-seq and RNA-seq and mapped endogenous G4 by ChIP-seq in wild type (WT) and BS cell lines. We observed that in BS cell lines, differential G4 formation positively correlated with both differential chromatin accessibility and gene expression. To test the direct involvement of G4s in the molecular phenotypes in BS, we applied pyridostatin, a G4-stabilizing molecule, in WT cells and showed that G4 stabilization partially recapitulated BS-associated molecular phenotypes. Additionally, we found that regions with increased chromatin accessibility in BS individuals in a family were also enriched for G4-forming sequences. Together, our data substantiate a regulatory role for G4s in Bloom syndrome and support a molecular model in which unresolved G4s in BLM- /- cells enhance chromatin accessibility, thereby promoting gene expression. These findings reveal an expanded regulatory function of BLM mediated through G4 structures and previously underappreciated role of G4s in the molecular etiology of BS.

## Introduction

Bloom Syndrome (BS, OMIM 210900) is a rare autosomal recessive cancer-predisposition disorder, first described in the 1950s ^1–3^. BS individuals manifest multifaceted clinical phenotypes. The prominent symptom is dwarfism due to pre-and postnatal developmental delay ^1,4^. Second, the affected individuals typically develop rashes on the face due to hypersensitivity to UV. They have increased risks for cancer and are prone to develop tumors at an early age, sometimes in their childhood. Additionally, they suffer from compromised immune function and are prone to infections. Thus, BS patients have a strikingly shorter lifespan, on average 26 years (see Cunniff et al. ^1^ for a detailed clinical summary).

In the 1990s it was discovered that BS is caused by loss-of-function mutations in the *BLM* gene, which encodes a 3′ to 5′ DNA helicase in the evolutionarily conserved RecQ helicase family ^1,2,5–7^. The BLM helicase resolves abnormal DNA structures—such as double-strand breaks, D-loops, forked duplexes, G-quadruplexes, and Holliday junctions—that arise during DNA replication, repair, and recombination ^8–10^. BLM therefore plays a crucial role in maintaining genome integrity.

Loss of BLM helicase function leads to profound alterations in cellular processes. At the molecular level, BS cells exhibit widespread dysregulations and features of genome instability. The hallmark and also diagnostic criterion of BS is excessive sister chromatid exchange events (SCEs), which occur at a rate seven-to ten-fold higher than in normal cells ^11,12^. BLM-deficient cells also display features underlying genome instability, impaired DNA repair, and elevated frequency of loss of heterozygosity, all of which increase the risk of developing cancer ^13–15^. These phenotypes reflect the critical function of BLM helicase in DNA repair, particularly its indispensable role in resolving double Holliday junctions without generating cross-over DNA products and thus suppressing illegitimate genetic recombination events ^16^. This latter feature of elevated recombination during mitosis has been leveraged to study and compare gene functions ^17,18^. Likely due to the lack of BLM to resolve aberrant DNA structures, BS cells exhibit lower DNA replication speed and have a slower cell cycle ^4,19^. Additionally, BS cells display altered gene expression profiles, and increased epigenetic age in terms of their DNA methylation patterns ^19–22^.

Some recent studies hinted at the relevance of G-quadruplexes (G4s) in the SCE events and transcriptomic changes in BS ^23–25,11^. G4s are four-stranded secondary DNA structures and a substrate for BLM helicase. They are formed by G-rich DNA sequences: four guanine bases pair with each other through a cyclic Hoogsten hydrogen bonding (as opposed to Watson–Crick G:C pairing) and form a square planar structure—G-quartet. Two or three G-quartets subsequently stack on top of each other to form a G4. Emerging studies have revealed that G4s regulate various processes such as DNA replication, transcription, and DNA-protein interactions ^26–30^. For instance, the formation of G4 could impede the progression of the DNA replication fork or machinery, which causes DNA damage and initiates DNA repair pathways ^31,32^. G4 structures have also been implicated in gene regulation. In transcription, the formation of G4 could impair the loading of RNA polymerase II, modulate the formation of R-loops, or stabilize the transcription bubble, thereby suppressing or enhancing transcription ^33,34^. G4s are prevalent in the promoter of many oncogenes, such as *c-MYC*, *c-KIT,* and *KRAS*, and it has been shown that their expression is influenced by the G4 forming status in their promoter. G4s are therefore considered a therapeutic target in cancer ^35–40^. Additionally, the formation of G4 can modulate DNA-protein interactions ^26,28,41^. For instance, G4 formation may recruit G4-binding proteins, displace transcription factors or even hinder histones binding to DNA, which could alter the local chromatin accessibility ^26,28,31,35^.

Due to the dynamic nature of G4 structures, their detection, especially in a cell-type specific manner, remains challenging ^27,42–44^. Bioinformatic tools have been often used to scan the DNA sequences and predict putative G4-forming sequences, known accordingly as G4 motifs ^27,45^. A given algorithm can detect 400,000 to over a million canonical G4 motifs (G_3+_N_1–7_G_3+_N_1–7_G_3+_N_1–7_G_3+_) *in silico* in the human genome ^46^. Another approach, G4-seq, measures polymerase stalling due to G4 *in vitro*, has reported over 500,000 G4-forming sequences in purified genomic DNA (herein referred to as G4-seq hits) ^43^. By contrast, G4 ChIP-seq or G4 CUT&Tag, which use an antibody–BG4–specifically targeting G4 structures, only identified 10,000 to 20,000 endogenous G4s in cells ^26,44,47–49^. The vastly different number of G4s identified via different methods imply that neither *in silico* G4 motifs nor *in vitro* G4-seq hits can reliably predict the G4 formation status in living cells. For this reason, we use “G4” in this article to specifically refer to actual secondary structures, as opposed to “G4 motifs” or “G4-seq hits”.

In the context of BS, recent studies showed the correlation between G4 motifs and certain molecular phenotypes ^11,21,24^. Van Wietmarschen et al. ^11^ mapped the locations of sister chromatid exchange events (the signature of BS) at single-cell resolution in cell lines derived from BS individuals and showed that sister chromatid exchange events exhibited a 1.0–1.2-fold enrichement at G4 motifs ^11^. Moreover, genes differentially expressed in BS cells are associated with the presence of G4 motifs ^21,24^. However, these analyses have been limited to G4 motifs rather than endogenous G4 structures. Given that G4 motifs vastly outnumber endogenous G4s and are invariant within a given genome, we reason that G4 motif alone is a poor predictor for G4 formation in living cells. Thus, it remains unclear if G4s are linked to the molecular pathology of BS. Particularly, it is as yet unresolved as to how the loss-of-function of BLM helicase influences endogenous G4 formation and—by extension—its subsequent impact on the molecular phenotypes observed in BS.

We thus set out to address this gap by firstly characterizing the molecular changes and secondly, determining if these changes are associated with endogenous G4 formation and the alterations in G4 dynamics. We mapped G4s and profiled chromatin accessibility and gene expression in cell lines derived from healthy donors (herein referred to as wild type or WT) and BS individuals. We found that changes in G4 formation in BS positively correlate with altered chromatin accessibility and gene expression. We further showed that G4 stabilization via pyridostatin (PDS) ^50–53^ can partially recapitulate molecular changes in BS. Analysis of a BS family further showed enrichment of G4-forming sequences in more accessible regions, further supporting the regulatory role of G4s in BS. To our knowledge, this is the first study to map endogenous G4 in BS and provides direct evidence that G4 could emerge as a central factor in the molecular etiology of BS and potentially serve as a therapeutic target. Our study further substantiates G4s’ regulatory function.

## Results

### BS cells exhibit disrupted molecular profiles

To characterize genome-wide molecular changes in BS, we collected two pairs of fibroblast and lymphoblastoid cell lines (LCL) derived from sex-matched and similarly aged BS and healthy donors (WT; see **Methods** for details) (**Fig. 1a**). We profiled the chromatin accessibility and transcriptome using ATAC-seq and RNA-seq, respectively. We also assayed endogenous G4 formation using G4 ChIP-seq ^47^, which utilizes an antibody specific to G4 structures and can directly map the G4s in cells as opposed to *in silico* G4 motifs or *in vitro* G4-seq hits from external sources to infer G4 formation status

**Figure 1.**
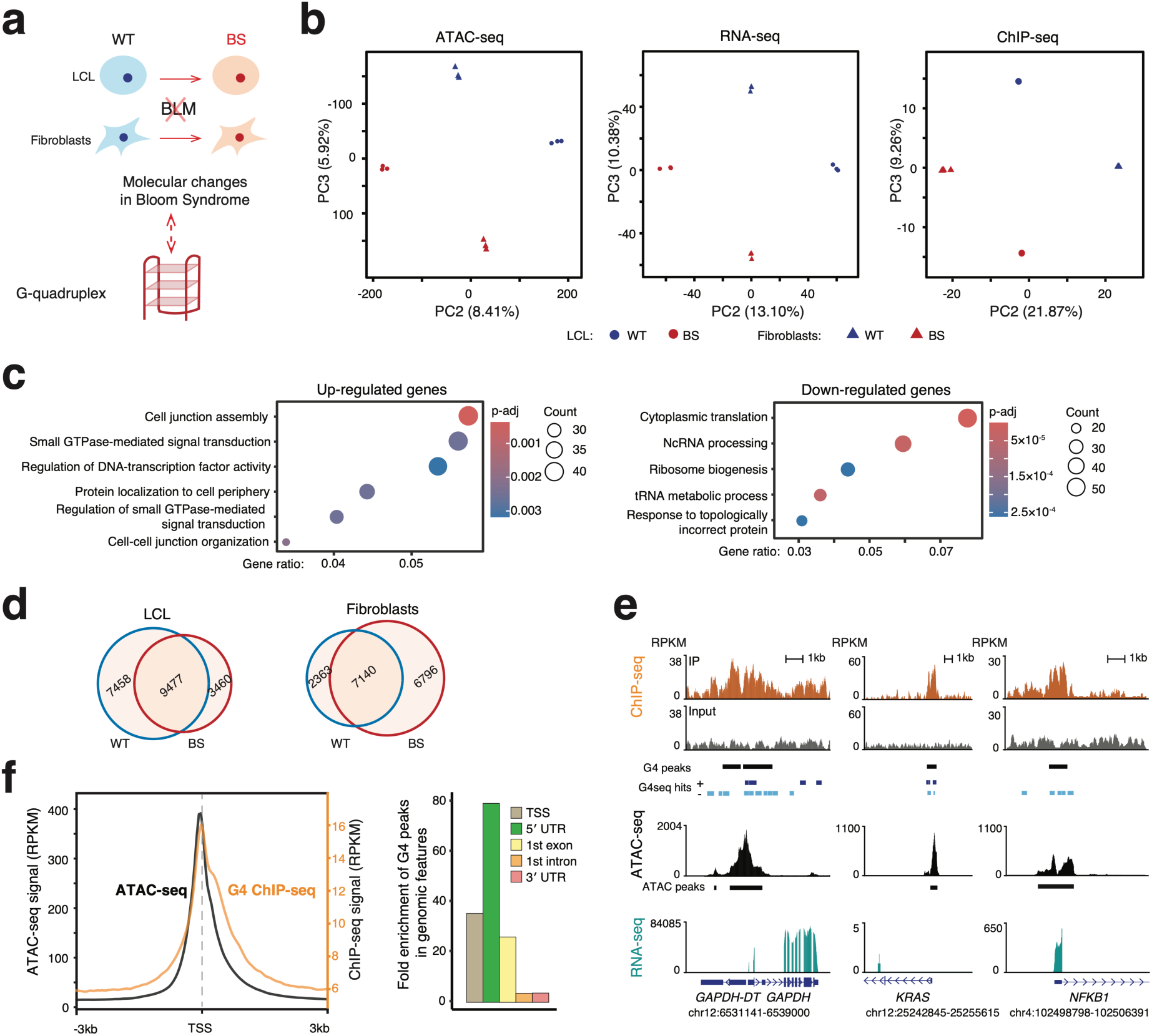
Bloom Syndrome samples exhibited disrupted molecular profiles. **(a)** Experimental design. LCL, lymphoblastoid cell lines; WT, wildtype; BS, Bloom Syndrome. **(b)** Principal component analysis of the genome-wide chromatin accessibility (left), gene expression profiles (middle) and endogenous G4 profile (right) in WT and BS samples. **(c)** Enriched GO pathways of shared up- and down-regulated genes in LCL-BS and fibroblast-BS. **(d)** Number of high-confidence G4 peaks in different samples. **(e)** Genome browser screenshots showing the G4 ChIP-seq, G4-seq hits, ATAC-seq and RNA-seq signals for *GAPDH* (housekeeping), *KRAS* (oncogene), and *NFKB1* (less-characterized G4-forming gene). **(f)** Distribution of G4 signals across genomic features. Left, averaged ATAC-seq and G4 ChIP-seq signal profiles within 6-kb regions centered on transcription start sites (TSSs). Right, fold enrichment of G4 peaks across different genomic features. Fold enrichment is calculated by permutation analysis (n = 1,000; *P* < 0.001 for all features; see **Methods**). UTR, untranslated region.

We first employed principal component analysis (PCA) to compare the global molecular profiles in WT lymphoblastoid (LCL–WT), BS lymphoblastoid (LCL–BS), fibroblast–WT, and fibroblast–BS cell lines. We observed in the PCA in each modality that the primary mode of variation (PC1; explaining over 70% variation across all three types of data) consistently separates samples by cell types, suggesting tissue-specific molecular profiles (**Suppl. Fig. 1a**). The next two major axes of variation, PC2 (8%, 13%, 22% for ATAC-seq, RNA-seq and ChIP-seq respectively) and PC3 (6%, 10%, 9% for ATAC-seq, RNA-seq and ChIP-seq respectively), separated the samples by disease status (BS vs. WT) in LCL and fibroblasts, respectively (**Fig. 1b**), implying distinct chromatin accessibility, gene expression and G4 profiles in BS and WT samples.

We next carried out differential analyses to characterize molecular signature in BS. Firstly, in ATAC-seq data, we identified on the order of 100,000 open chromatin regions (101,787, 115,027, 125,352, and 114,427 in LCL-WT, LCL-BS, fibroblast-WT, and fibroblast-BS cells, respectively) and detected the differential signals with *DESeq2* ^54^. Of note, in fibroblast samples, to mitigate the impact of copy number differences on differential analysis, we developed and applied a pipeline featuring copy number normalization (see **Methods**) ^55^. In the LCL-BS vs. LCL-WT comparison, we identified 72,580 (57%) significantly differentially accessible (DA) chromatin regions (adjusted p-value (p-adj) < 0.05), of which 39,920 and 32,660 displayed increased (BS > WT) and decreased (BS < WT) chromatin accessibility, respectively (**Suppl. Fig 1b**, **left**). In the fibroblast-BS vs. fibroblast-WT comparison, we identified a similar number (78,789, or 60%, CNV-corrected, of which 36,225 (25%) and 42,564 (35%) were significantly more or less accessible, respectively) ^55^ (**Suppl. Fig. 1b**, **right)**. Notably, despite about 8,000 shared DA peaks, LCL-BS showed a weak anti-correlation vs. and fibroblast-BS peaks (Pearson correlation, r = −0.11, *P* < 0.001; **Suppl. Fig. 1c**), implying the impacts of BS on the chromatin states can and do show cell-type specificity.

Next, we characterized gene expression changes in BS vs. WT for each cell type. Considering genes expressed in at least one sample, we identified 6,703 significantly differentially expressed (DE) genes (p-adj < 0.05; 3,627 up-vs. 3,076 down-regulated) in LCL and 11,140 in Fib (p-adj < 0.05; 5,347 up-vs. 5,793 down-regulated) (**Suppl. Fig. 1d**). Consistent with the cell-specific changes in BS in the PCA, the DE gene sets in LCL and Fib were distinct (1,911 in the same direction vs. 1,736 in opposite directions) and displayed significant, but very weak anti-correlation (Pearson correlation, *r* = −0.03, *P* < 0.001; **Suppl. Fig. 1e**).

To determine if specific molecular functions were enriched among differentially expressed (DE) genes, we conducted a Gene Ontology (GO) analysis using DE genes ranked by fold changes. We first focused on DE genes shared across both cell types to identify commonly affected molecular functions in BS. Up-regulated genes in BS were enriched for DNA-transcription factor activity, cell junction, cell adhesion, and signaling transduction (**Fig.1c**, **left**). Down-regulated genes were associated with translation, ribosome metabolism, and ncRNA processing (**Fig.1c, right**). Notably, these categories correspond to previous study that BLM can unwind G4 and facilitate RNA polymerase I-mediated ribosomal RNA transcription ^56,57^. Another notable enriched pathway across both cell types is the hallmark gene set of MYC targets, which is closely related to tumorigenesis (**Suppl. Fig. 1f**; **Suppl. Fig. 1g**). Moreover, *MYC* is a well-known oncogene whose expression level is highly dependent on the formation of G4 in its promoter ^35,36^.

Within cell types (LCL and fibroblasts), we found down-regulated genes in BS–LCL to be associated with immune functions such as the “activation of the immune responses” and “immunoglobulin production”, implying potentially decreased immune functions in BS, a finding reminiscent of the immunodeficiency in BS individuals (**Suppl. Fig. 1f**) ^1^. In BS–Fib, they were mostly related to cell skeleton organization and cell morphology-related structure development, such as “muscle structure development” (**Suppl. Fig. 1g**). These pathways provide insights into the developmental delay or smaller body size in BS individuals.

As a third functional channel in our data, we profiled the endogenous G4s in WT and BS cells via G4 ChIP-seq ^47^. Here, we used *MACS2* ^58^ to call G4 peaks and applied the irreproducible rate (IDR) of 0.05 to filter the reproducible peaks across biological replicates via *idr* ^59^. We identified on average about 14,000 high-confidence endogenous G4s (10,382, 15,204, 17,125, and 13,415 in LCL-WT, LCL-BS, fibroblast-WT and fibroblast-BS cells, respectively). Among these, 4,977 G4 sites were shared across all samples (corresponding to 44.8%, 35.8%, 29.4% and 38.4% in LCL-WT, LCL-BS, fibroblast-WT, fibroblast-BS, respectively) were shared across all samples, indicating that there are many cell type and disease-status specific G4 sites (**Fig. 1d**; **Suppl. Fig. 2a**). Motif analyses via *MEME* ^60^ on these G4 peaks confirmed strong G4-forming sequence signatures of the triple and/or quadruple Guanine-repeats (**Suppl. Fig. 2b**, e-value < 1 × 10^−15^ for all motifs shown). Furthermore, about 80% of these detected G4 peaks overlapped with known G4-seq hits (**Suppl. Fig. 2c**) ^31^.

A number of previous studies have reported that endogenous G4s are prevalent in the promoter of highly expressed housekeeping genes and some oncogenes ^26,29,36,37,39,47^. Here, we detected G4 peaks in the housekeeping genes such as *GAPDH* and *H3*, in oncogenes such as *KRAS*, *MYC*, and *KIT*. We also detected G4s in *NFKB1*, a less well-characterized and LCL-specific G4-forming gene, *NFKB1* (**Fig. 1e**). These peaks coincided with open-chromatin signatures (**Fig. 1e**). Globally, 70% ∼ 80% of G4 peaks overlapped with ATAC-seq peaks (**Suppl. Fig. 2d**). Particularly in the promoter regions, G4s and ATAC-seq displayed highly similar signal profiles with both signals peaking slightly 5′ to the TSS (**Fig. 1f**). These observations are mechanistically intuitive, as G4s form from single-stranded DNA and are expected to be found in open chromatin regions, especially at promoters where active transcription generates transient single-stranded DNA ^26,44^.

In terms of gene features, G4 signals are highly abundant in the gene body, particularly near the TSS with about 80% of G4 peaks residing in the promoters (defined as TSS ± 3kb); with over 50% directly overlapping a TSS (**Suppl. Fig. 2e**, **2f**). Moreover, G4 peaks exhibited a significant enrichment in the 5′ UTR, TSS, first exon, first intron, first exon-intron (1st-Ex-Int) junction, and 3′ UTR compared to randomly shuffled G4 peaks regions of the same length (permutation, n = 1000; *P* < 0.001 for all features; see **Methods**) (**Fig. 1f**; **Suppl. Fig. 2g**).

Building on this and given that previous studies reported conflicting results on the enrichment of G4 forming sequences at the 1st-Ex-Int junctions of DE genes in BS ^24,25^, we revisited this question in each cell type using our endogenous G4 sites. We first assessed the enrichment at all expressed genes and then compared the enrichment in DE genes relative to this background. While there is a background enrichment in both cell types, we only observed significant enrichment of G4 peaks at the 1st-Ex-Int junctions of DE genes in fibroblasts, but not in LCLs (permutation, n = 1000; fibroblast, *P* < 0.001; LCL: *P* = 0.15; see **Methods**) (**Suppl. Fig. 3a-d**). This result is notable, in that it shows that the 1st-Ex-Int enrichment may itself be cell-type-specific rather than constitutive. Importantly, this discrepancy highlights the distinction between putative G4-forming sequences and endogenous G4s, underscoring the value of directly mapping endogenous G4s to investigate their functional relevance.

To address the role of G4 specifically in the molecular phenotypes of BS, we characterized the differential G4-forming sites (DF G4 sites) in BS vs. WT within each cell type with *DiffBind* ^61,62^, contrasting BS vs. WT cells within each cell type. In LCL-BS vs. LCL-WT, we identified 2,436 (13.79%) significantly DF G4 sites (FDR < 0.05), 1,401, and 1,035 with increased and decreased G4 formation, respectively (**Suppl. Fig. 1h**). In fibroblasts samples, we detected 2,022 (10.07%) and 1,764 (8.49%) sites with increased and decreased G4 formation (after normalized for copy number as in ATAC-seq ^63^; **Suppl. Fig. 1h**).

Together, we observed that G4s were predominantly localized to functional genomic elements, particularly promoter regions and open chromatin, consistent with their proposed regulatory roles and supporting the quality of our G4 ChIP-seq data. Across all data modalities, BS cells exhibited disrupted molecular profiles above and beyond cell-type specific effects. In particular, BS cells displayed a distinct G4-formation landscape. Given the enrichment of G4s at regulatory elements, alterations in G4 dynamics are likely to have downstream molecular consequences in chromatin accessibility and gene expression.

### G4 formation changes positively correlate with chromatin accessibility changes in Bloom Syndrome cells

We next investigated how the different regulatory modalities may affect one another, namely between ATAC-seq, RNA-seq and G4 ChIP-seq. We first looked at the DA peaks and DE genes. To do so, we linked an ATAC-seq peak to the potential target gene by proximity to the nearest TSS (+/-3kb). In both cell types, we observed a strong positive correlation between DA peaks in promoters and DE genes (Pearson correlation; LCL, *r* = 0.58, *P* < 2.2 × 10^−16^; Fib, *r* = 0.53, *P* < 2.2 × 10^−16^) (**Fig. 2a**; **Suppl. Fig. 4a)**. This is in line with previous studies that chromatin accessibility changes in promoter regions largely correlated with gene expression changes ^64–66^.

**Figure 2.**
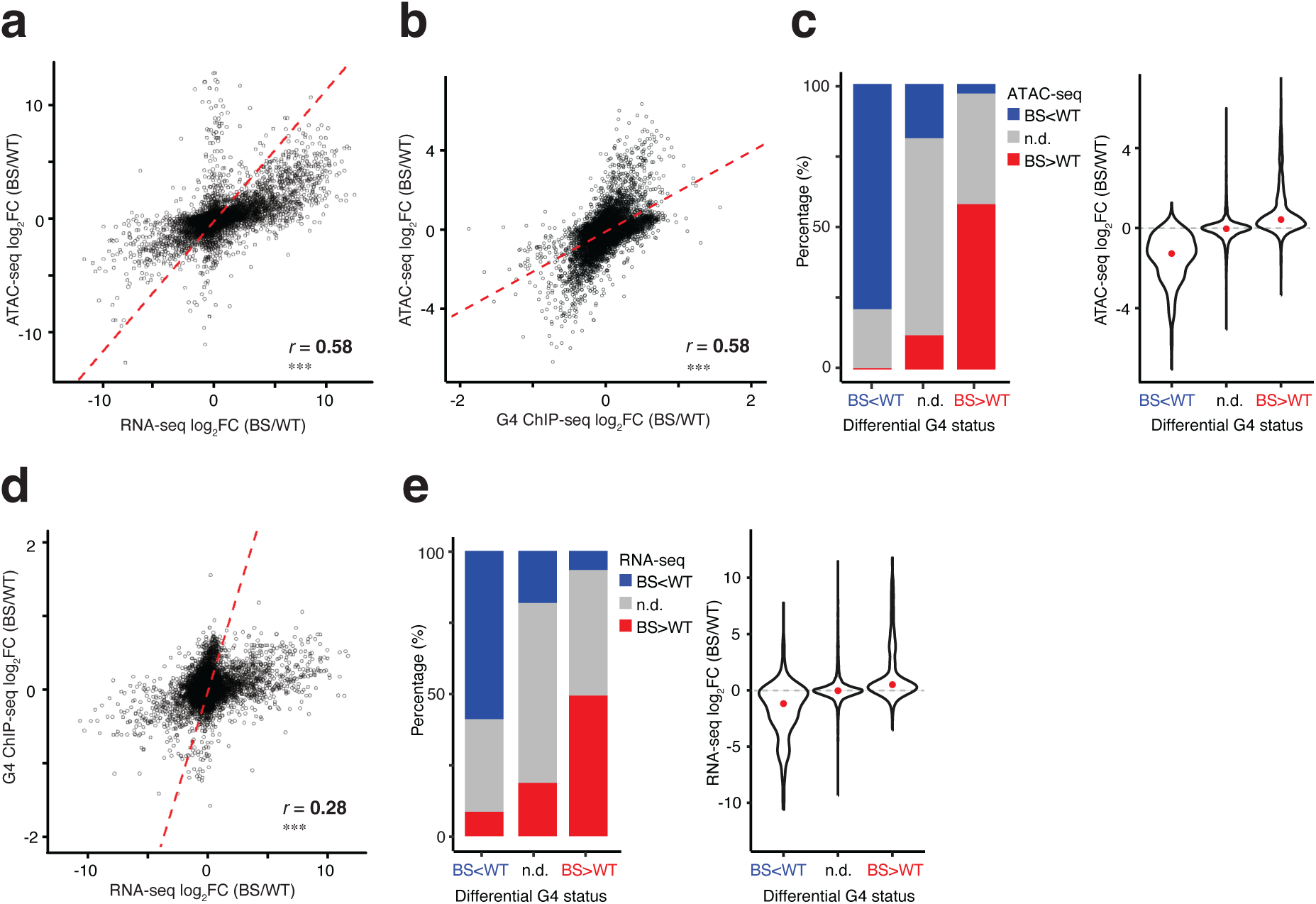
Differential G4 formation positively correlated with molecular phenotypes in Bloom Syndrome. **(a)** Positive correlation between differential chromatin accessibility and differential gene expression in LCL-BS vs. LCL-WT. The red line indicates the fitted linear regression. R is the Pearson correlation coefficient. ***, *P* < 2 × 10^-16^. **(b)** Positive correlation between differential chromatin accessibility and differential G4 formation in LCL-BS vs. LCL-WT. The red line indicates the fitted linear regression. R is the Pearson correlation coefficient and ***, *P* < 2 × 10^-16^. **(c)** Distribution of differentially accessible chromatin regions and their fold changes stratified by changes in G4 formation in LCL-BS vs. LCL-WT. Red dots denote the median log_2_FC. N.d., non-differential signals with p-adj ≥ 0.05. **(d)** Positive correlation between differential gene expression and differential G4 formation in LCL-BS vs. LCL-WT. The red line indicates the fitted linear regression. R is the Pearson correlation coefficient. ***, *P* < 2 × 10^-16^. **(e)** Distribution of differentially expressed genes and their fold changes stratified by the changes in G4 formation in LCL-BS vs. LCL-WT. Red dots denote the median of log_2_FC.

To examine the dynamics between chromatin accessibility and G4 formation, we intersected the ATAC-seq peaks and G4 sites for each cell line. We identified in total 12,680 and 16,345 overlapping ATAC–G4 peaks in LCL and fibroblast, respectively. Overall, we also found significant positive correlation among changes in these two data modalities (Pearson correlation; LCL, *r* = 0.58, *P* < 2.2 × 10^−16^; fibroblasts, *r* = 0.59, *P* < 2.2 × 10^−16^) (**Fig. 2b**; **Suppl. Fig. 4b**). As G4 ChIP-seq signals progressively shifted from decreased to increased formation in BS, chromatin accessibility exhibited concordant directional changes (**Fig. 2c**; **Suppl. Fig. 4c**). Specifically, 79% of regions with decreased G4 signals in BS overlapped open chromatin regions with decreased accessibility, whereas 57.9% of regions with increased G4 signals overlapped regions with increased chromatin accessibility (**Fig. 2c**; **Suppl. Fig. 4c**). Conversely, only 3.4% of regions with increased G4 formation were associated with decreased chromatin accessibility (**Fig. 2c**; **Suppl. Fig. 4c)**.

These observations revealed a strong positive concordance between changes in G4 formation and chromatin accessibility in BS cells.

### G4 formation changes positively correlated with gene expression changes in Bloom Syndrome cells

Given the proposed role of G4 in regulating gene expression, we next asked whether changes in G4 formation are associated with gene expression changes. To investigate this, we assigned a G4 peak to a gene if it was located within the gene’s promoter, yielding 9,272 and 11,657 G4–gene pairs in LCL and fibroblasts, respectively. We found that differential gene expression correlates positively with differential G4 formation, with fibroblasts showing a stronger correlation (Pearson correlation; *r* = 0.28 and 0.37 in LCL and fibroblasts, both *P* < 2.2 × 10^−16^; **Fig. 2d**; **Suppl. Fig. 4d**). Similarly, stratifying G4–gene pairs by the directionality of DF sites, we observed that the majority of changes in G4 levels were consistent with changes in gene expression. Genes with significantly increased G4 formation in the promoter mostly exhibited increased expression (49.3%) and conversely, genes with decreased G4 formation were mainly downregulated (59.1%) (**Fig. 2e**; **Suppl. Fig. 4e**). Across both cell types, gene expression changes in BS were positively associated with endogenous G4 dynamics, with increased promoter G4 formation linked to higher gene expression.

Together, the coordinated changes in G4 formation, chromatin accessibility, and gene expression suggest that G4s may represent an underappreciated regulatory layer in BS.

### PDS treatment partially recapitulates the molecular changes in Bloom Syndrome cells

Pyridostatin (PDS) is a G4-stabilizing small molecule^50–52^. It intercalates into G4 and increases the required physical force to unfold G4, thereby inhibiting G4 resolution via BLM ^53^. To further investigate the causal role of G4, we thus applied PDS to WT-LCL and fibroblasts cell lines to mimic the defective G4-resolving abilities in BS. Following titration experiments to determine the optimal PDS concentrations, we treated WT cells for 24h with 10mM PDS (vs. a DMSO carrier, herein referred as control) and then profiled their respective open chromatin landscape and transcriptome (**Fig. 3a**)

**Figure 3.**
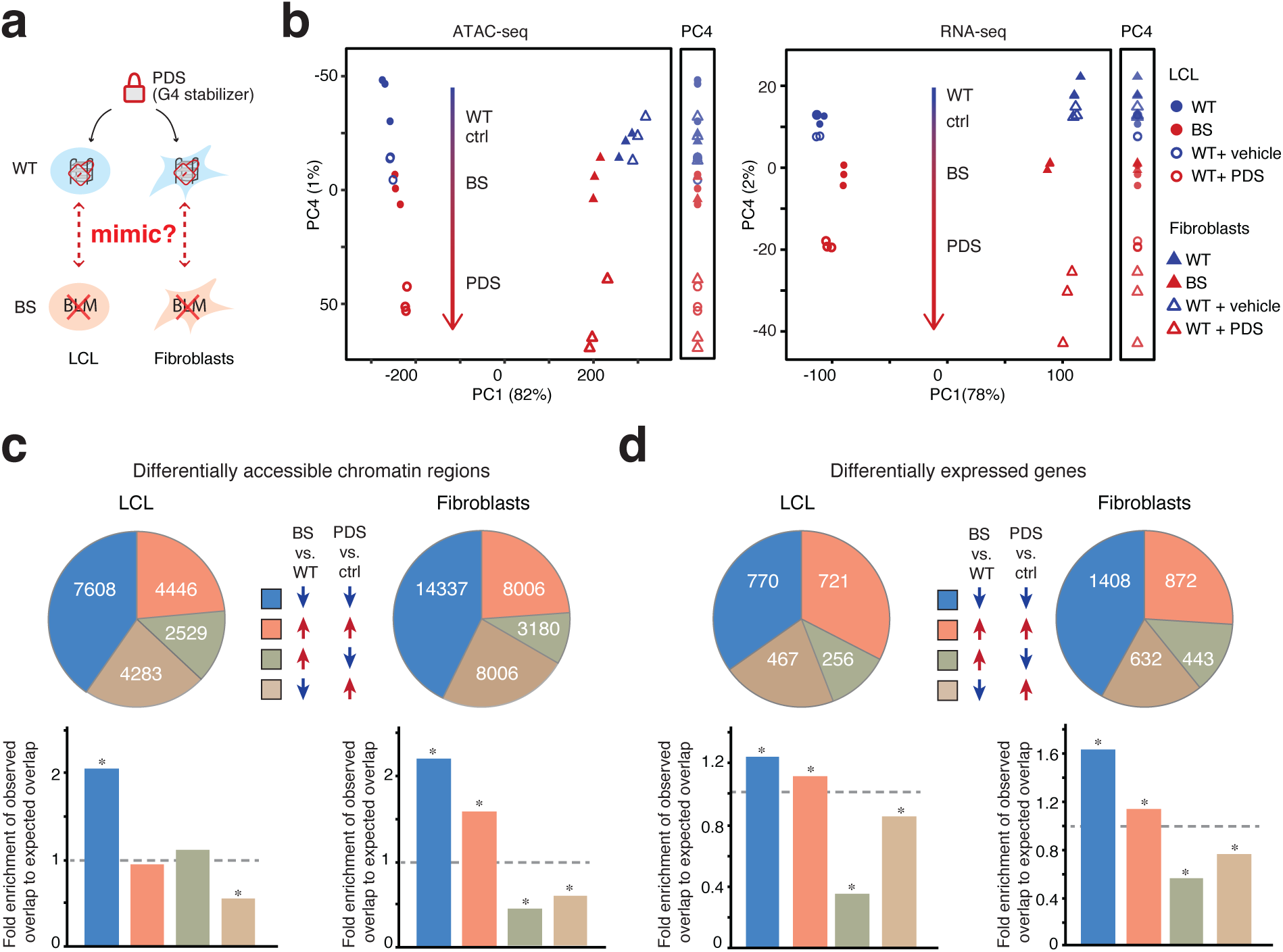
G-quadruplex stabilization via PDS partially recapitulated chromatin accessibility and gene expression changes in Bloom Syndrome. **(a)** Experimental design. PDS, pyridostatin, a G4 stabilizing molecule. **(b)** Principal component analysis of the genome-wide chromatin accessibility (left) and gene expression profiles (right) in samples across all cell types and conditions. **(c)** Overlap of differentially accessible chromatin regions (top) and their fold enrichment relative to expected (bottom) in Bloom Syndrome and under PDS treatment in LCL and Fib, respectively. *, hypergeometric tests, *P* < 0.001 (**Suppl. Table 2**). **(d)** Overlap of differentially expressed genes (top) and fold enrichment relative to random expectation (bottom) in Bloom Syndrome and under PDS treatment in LCL and fibroblast cell lines, respectively. *, hypergeometric tests, *P* < 0.001 (**Suppl. Table 3**). WT samples treated with vehicle are referred to as the control condition (Ctrl).

We performed PCA to compare the global ATAC-seq profiles across all samples, including both PDS and BS contrasts. PCA revealed distinct chromatin accessibility profiles among samples of different tissue origin, with PC1 explaining approximately 80% of the variance (**Fig. 3b**, **left**). Notably, we observed that samples fall along a spectrum on PC4 that separated samples expected to have normal vs. elevated G4 levels, namely WT, control, BS and finally PDS–treated cells (**Fig. 3b**, **left**). This axis highlights the common impacts between Bloom Syndrome and PDS treatment on chromatin accessibility regardless of tissue origin. PC2 (7%) and PC3 (6%) separated Bloom Syndrome samples from the remaining samples in LCL and fibroblasts, respectively, reflecting more cell-specific effects (**Suppl. Fig. 5a**). We next carried out PCA on gene expression profiles across all the samples and observed comparable results. PC1 (78%) again separated samples by cell types (**Fig. 3b**, **right**) while PC2 (10%) and PC3 (7%) represented the impacts of BS in LCL and fibroblasts, respectively. PC4 recapitulated the same WT-control-BS–PDS spectrum, suggesting a shared transcriptomic signature in BS and PDS–treated cells (**Fig. 3b**, **right**).

Next, we characterized the differential signals. In LCL, PDS treatment resulted in 17,154 and 16,580 peaks with significantly increased and decreased signals, respectively (combined: 33,734, or 27% of all peaks **Suppl. Fig. 5b**, **left**). In fibroblasts, we detected 42,301 (32%) DA peaks, comprising 20,210 with increased accessibility and 22,091 with decreased accessibility (**Suppl. Fig. 5b**, **right**).

To assess whether PDS treatment recapitulates those changes in chromatin accessibility in BS, we compared differential signals caused by BS and upon PDS treatment in each cell type and observed a significant positive correlation in both (Pearson correlation; LCL, r = 0.29, *P* < 2.2 × 10^−16^; fibroblasts, r = 0.44, *P* < 2.2 × 10^−16^). For convenience, we use the notations BS+, BS–, PDS+ and PDS– to refer to changes relative to their control. In LCL, we found 4,446 BS+/PDS+ (23.6% of peaks differentially accessible in both conditions; percentages in the rest of this are calculated the same way) and 7,608 BS–/PDS– regions (40.3%) (**Fig. 3c**, **top-left**). In fibroblasts, we found 9,060 BS+/PDS+ (28.7%) and 15,036 BS–/PDS– regions (50.8%) (**Fig. 3c**, **top-right**). In both cell types, the observed numbers of BS–/PDS– peaks were significantly higher than expected, with approximately two-fold enrichment (hypergeometric test; Fib, *P _(> observed)_* = 1.6 × 10^−6^; LCL, *P _(> observed)_ =* 0.003) (**Fig. 3c**, **bottom**). BS+/PDS+ is also significantly enriched, but only in fibroblasts and not LCL (**Fig. 3c**; hypergeometric test; fibroblasts, *P _(> observed)_* < 1×10^−16^; LCL, *P _(> observed)_* = 1.00). In contrast, there were less than expected BS–/PDS+ peaks in both cell types and a significant depletion of BS+/PDS– peaks in Fib (**Fig. 3c**; hypergeometric test; *P* _(< observed)_ in **Suppl. Table 2**).

Next, we compared RNA-seq data upon PDS or DMSO treatment and identified 6,258 (39%) significantly DE genes in LCL (2,930 upregulated and 3,328 downregulated) and 4,828 DE genes (2,363 upregulated and 2,465 downregulated) in fibroblasts (**Suppl. Fig. 5d)**. Importantly, in both cases, the DE genes showed a strong correlation with those identified in the corresponding BS cell line of the same cell type (Pearson correlation; r = 0.29 and 0.44 in LCL and fibroblasts, both *P* < 2.2 × 10^−16^). Similar to DA peaks, the number of BS+/PDS+ (LCL, 32.6%; Fib, 26.0%;percentages are relative to the total number of differentially expressed peaks in both conditions) and BS–/PDS– (LCL, 34.8%; Fib, 42.0%) DE genes was significantly enriched in both cell types, displaying a 1.2 to 2-fold enrichment (**Fig. 3d**, **bottom**; hypergeometric test; *P _(> observed)_* in **Suppl. Table 3**). Conversely, there was a depletion of BS–/PDS+ and BS+/PDS– genes (**Fig. 3d**, **bottom**; hypergeometric test; *P _(< observed)_* in **Suppl. Table 3**).

In contrast to the more cell type-specific effects of BS (**Suppl. Fig. 1e**; **Suppl. Fig. 5a**), we also noticed that DA peaks and DE genes induced by PDS treatment exhibited stronger correlation across cell types (Pearson correlation for differential signals in LCL and in fibroblasts upon PDS treatment and in BS; ; DA peaks: r = 0.37 vs. −0.11; DE genes: r = 0.84 vs. 0.07; both *P* < 2.2×10^−16^) and smaller fold changes compared to those in BS cells (**Suppl. Fig. 5c**, **5e**). These suggest that PDS treatment elicited more universal but less extensive responses on both chromatin accessibility and gene expression.

To summarize, we observed that PDS treatment can partially recapitulate both the global and differential chromatin accessibility and gene expression features caused by BS. This indicates potential shared regulatory mechanisms between G4 stabilization and BS, highlighting the role of G4 in a suite of BS molecular phenotypes that may have previously been understood to have other causes. Moreover, chromatin accessibility changes in response to PDS treatment imply a possible feedback mechanism between G4 structure and nucleosome positioning.

### G4 formation potentiates gene expression by increasing chromatin accessibility

Having established a strong association between G4 dynamics and molecular changes in BS, we next sought to investigate the molecular mechanisms linking G4 formation to chromatin accessibility and gene expression.

To examine whether G4 formation may affect chromatin accessibility, we stratified the ATAC-seq peaks based on their overlap with G4 peaks (G4+ ATAC peaks and G4– ATAC peaks). We observed that G4+ ATAC-seq peaks are wider in LCL-WT (median size: 988bp vs. 347bp; Wilcoxon test, *P* < 2 × 10^−16^) and exhibited overall higher chromatin accessibility (median FPKM: 31.1 vs. 5.96; Wilcoxon test, *P* < 2 × 10^−16^; **Fig. 4a**; for brevity we report results from LCL-WT and we observed similar effects in the fibroblasts). Additionally, we observed a higher density of open chromatin regions around G4 sites (Wilcoxon test, *P* < 2 × 10^−16^, **Fig. 4b**; **Suppl. Fig. 6a**), implying G4 formation activities may not only influence the overlapping ATAC-seq peaks but also proximal regions.

**Figure 4.**
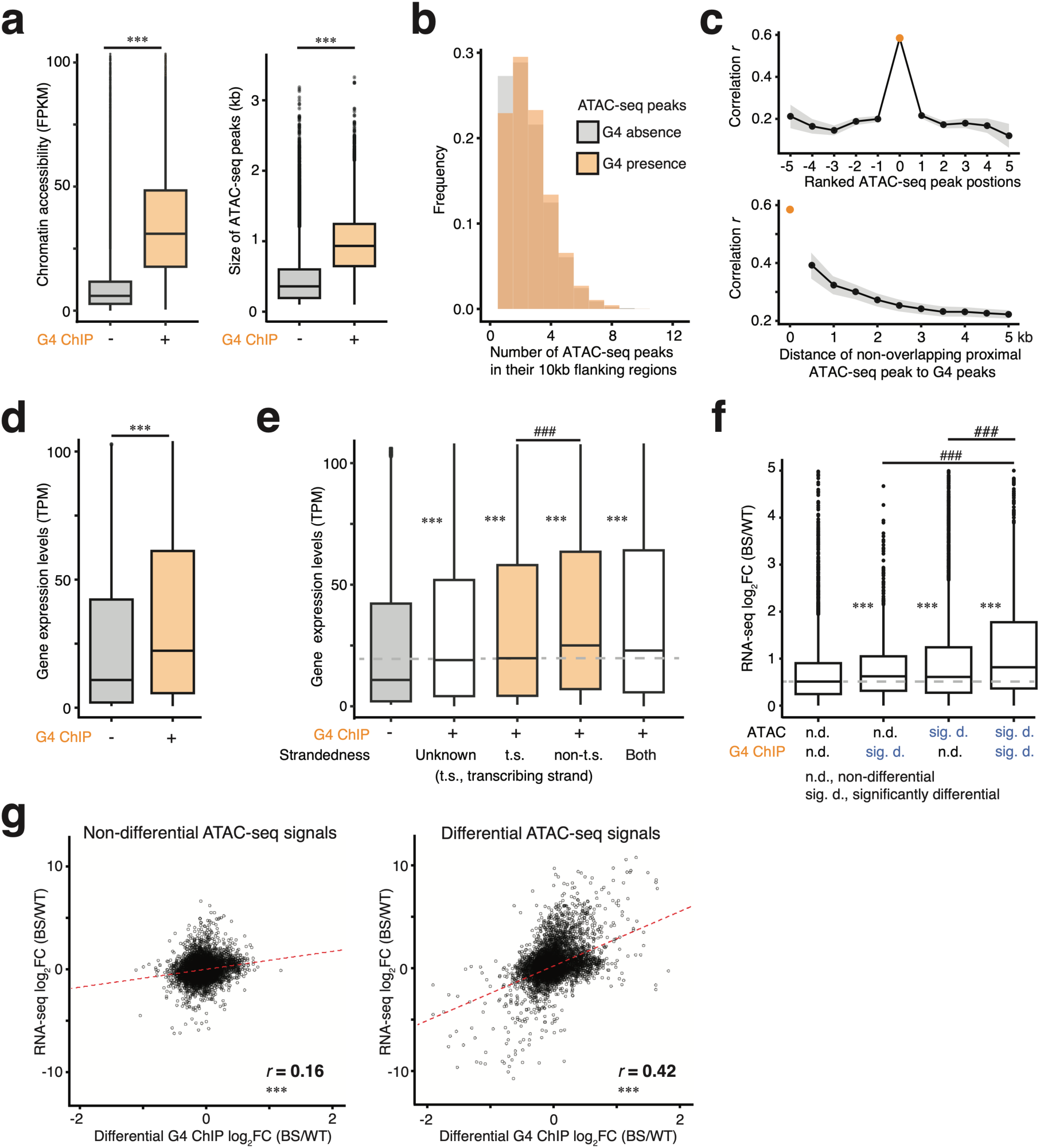
Molecular insight into regulatory functions of G-quadruplexes in Bloom Syndrome. **(a)** ATAC-seq peaks overlapping G4 peaks exhibit higher chromatin accessibility (left) and broader open chromatin regions (right). **(b)** The presence of G4 peaks is associated with increased local open chromatin density. (**c**) Correlation between G4 ChIP-seq log₂ fold change (log₂FC) and ATAC-seq log₂FC for G4-overlapping and G4-proximal ATAC-seq peaks. A strong positive correlation is observed for G4-overlapping open chromatin regions, which is markedly reduced for G4-proximal but non-overlapping regions. **(d)** Genes with G4s in the promoter exhibit higher gene expression. **(e)** Genes with G4 on the non-transcribing strand show higher expression than those with G4 on the transcribing strand. Wilcoxon tests were used. Groups marked with * were compared to genes without G4s (leftmost boxplot); ***, *P* < 2 × 10^-16^; ###, *P* = 8 × 10^-4^. **(f)** Additive effects of differential chromatin accessibility and G4 formation on differential gene expression. Wilcoxon tests were used. Groups marked with * were compared to genes without significant changes in chromatin accessibility or G4 formation (leftmost boxplot); **, *P* = 0.003; ***, *P* < 2 × 10^-16^; ###, *P* < 2 × 10^-16^. **(g)** Correlation between differential G4 formation and differential gene expression, stratified by significant changes in chromatin accessibility. The red line denotes the fitted linear regression. R is the Pearson correlation coefficient and ***, *P* < 2 × 10^-16^.

We further evaluated if the afore-described positive correlation between DA peaks and DF G4 sites extended to the proximal ATAC-seq peaks besides the overlapping ones. The proximity of ATAC-seq peaks was assessed both by their relative ranked position to G4 sites (0 for overlapping, positive for upstream, and negative for downstream; **Suppl. Fig. 6b**) or physical distance. We observed that correlation was strongest with directly overlapping ATAC-seq peaks and decayed sharply as the ranking or physical distance increased (ranking: **Fig. 4c, top**; **Supp. Fig. 7a**, **top**; physical: **Fig. 4c**, **bottom**; **Supp. Fig. 7a, bottom**). To summarize, we observed that G4 formation coincides with broader and more accessible chromatin regions. Combined with the positive correlation between G4 changes and chromatin accessibility—particularly at G4-overlapping ATAC peaks rather than adjacent regions—these findings strongly suggest that once formed, G4s can feed back to reinforce chromatin accessibility, predominantly in the regions where G4 forms. Consequently, any changes in G4 formation are likely to subsequently influence chromatin accessibility. At the same time, we acknowledge that this does not by itself establish a hierarchy between G4 formation and open chromatin, since G4 structures arise in single-stranded DNA and thus require open chromatin.

To investigate how G4 formation influences gene expression, we similarly stratified the expressed genes based on the presence or absence of G4 peaks in their promoters (G4+ genes and G4− genes). Similar to the ATAC-seq analysis, G4+ genes showed significantly higher expression levels (median TPM: 22.2 vs. 10.8; Wilcoxon test, *P* < 2 × 10^−16^) (**Fig. 4d**). Moreover, the average gene expression tends to decrease with increasing distance from the G4 peak to the TSS (**Suppl. Fig. 6c**). Next, given that one DNA strand is transcribed, G4s may influence gene expression differently depending on which strand they form ^33^. To examine this, we intersected our unstranded G4 peaks with the publicly available strand-resolved G4-seq data ^26^ and used this to infer their strandedness if a G4 peak overlaps only G4-seq hits from the same DNA strands (**Suppl. Fig. 6d**). G4 peaks overlapping no G4-seq hit or multiple G4-seq hits from both DNA strands can be not inferred. Ultimately, we were able to assign strandedness to 50% of G4 peaks (**Suppl. Fig. 6e**). We found that regardless of their strandedness, genes with G4s showed higher expression (**Fig. 4e**, Wilcoxon test, *P* < 2 × 10^−16^). Particularly, genes with G4s on the non-transcribing strand showed even higher gene expression than genes with G4s on the transcribing strand (**Fig. 4e**, Wilcoxon test, *P* = 8 × 10^−4^). These observations showed that G4 formation promotes gene expression, especially when G4s form on the non-transcribing strand. This could explain that in BS vs. WT, genes with increased G4 formation mostly showed increased gene expression.

Next, given the three-way correlation between ATAC-seq, G4 ChIP-seq and RNA-seq (**Fig. 2a**, **2b**, **2d**), we investigated whether G4 changes and chromatin accessibility changes may act cooperatively on gene expression. To address this, we focused primarily on the subset of genes that were expressed and overlapped with both G4 and ATAC-seq peaks—identifying 9,194 and 11,580 such genes in LCL and fibroblasts, respectively. Stratifying genes by the differential status of their associated ATAC-seq and G4 peaks and compared the strength of gene expression changes, we found that genes with a significant change in either modality showed increased expression (Wilcoxon test; *, *P* < 2 × 10^−16^) (**Fig. 4f, Suppl. Fig. 7b**). Importantly, genes with significant changes in both chromatin accessibility and G4 formation showed the strongest upregulation, suggesting additive effects (Wilcoxon test; #, *P* < 2 × 10^−16^) (**Fig. 4f**; **Suppl. Fig. 7b**). To further assess whether there is a hierarchy in the regulation of gene expression by G4 and chromatin accessibility, we analyzed gene expression correlations stratified by the differential status of the third data modality. Interestingly, G4 changes had minimal impact on the positive correlation between chromatin accessibility and gene expression, whereas the correlation between G4 changes and gene expression was largely dependent on significant changes in chromatin accessibility (**Fig. 4g**; **Suppl. Fig. 6h**; **Suppl. Fig. 7c, 7d**). These results suggest that altered G-quadruplexes regulate gene expression primarily by modulating chromatin accessibility, rather than acting independently or through an alternative regulatory hierarchy. Importantly, this also points to a reciprocal interaction between G4 formation and chromatin accessibility.

To formally test this feedback mechanism wherein G4 formation promotes the further opening of chromatin, we built a multifactor ANOVA model (RNA = ATAC + G4 + ATAC × G4), modeling gene expression changes as a function of the differential status of chromatin accessibility and G4 formation. The analysis revealed a significant interaction between chromatin accessibility and G4 formation (Type III ANOVA; F = 3.89, *P* _(> F)_ = 0.004), supporting a cooperative model. It also confirmed that both differential ATAC-seq and differential G4 formation status imposed significant impacts on gene expression changes (Type III ANOVA; ChIP: F = 184.2, *P* _(> F)_ < 2 × 10^−16^; ATAC: F = 158.11, *P* _(> F)_ < 2 × 10^−16^).

Taken together, G4 formation and open chromatin status are both associated with enhanced gene expression, and they could act cooperatively to modulate transcriptional output. Our results further imply the hierarchy that G4s likely exert their regulatory effects by promoting chromatin accessibility. This also supports our reciprocal modulation model between G4 and chromatin accessibility in which G4 formation requires open chromatin environment and once formed, G4s reinforce chromatin accessibility.

### Differentially accessible chromatin regions in Bloom Syndrome individuals are enriched for G4-seq hits

To verify our results, we acquired lymphocytes isolated from a Bloom Syndrome family with the BS-causing Ashkenazi mutation (*blm*^Ash^), *BLM* c.2207-2212delATCTGAinsTAGATTC from the Bloom Syndrome Registry (**Fig. 5a**; **Suppl. Table 4**) ^7^. Due to the limited material, we were only able to assay the open chromatin landscape via ATAC-seq and data from the carrier mother and carrier daughter were excluded due to low data quality. In total, we identified 60,913 peaks from non-BS and BS individuals and 982 significantly differentially accessible chromatin regions (p-adj < 0.05; 60 with increased signal and 922 with decreased signals) (**Suppl. Fig. 8**). To determine if the differentially accessible chromatin regions are associated with G4, we used *in vitro* G4-seq hits as a proxy for G4 due to the lack of corresponding endogenous G4 data for these samples. Specifically, we estimated the enrichment of G4-seq hits in ATAC-seq peaks by comparing the observed overlap to the expected overlap estimated by permutation, in which we counted the number of G4-seq hits in open chromatin regions randomly shuffled in the genome (see **Methods**). Overall, open chromatin regions were enriched for G4-seq hits (permutation, n=1000; observed overlap: 38514; expected overlap: 8290 ± 108.8; z-test, *P* < 0.001; fold enrichment = 4.65). Interestingly, the more accessible chromatin regions in BS individual (permutation, n=1000; observed overlap: 24; expected overlap: 10.5 ± 3.81; z-test, *P* < 0.001; fold enrichment = 2.29) exhibited higher enrichment for G4-seq hits compared to those less accessible chromatin regions (permutation, n=1000; observed overlap: 85; expected overlap: 65.3 ± 8.55; z-test, *P* < 0.01; fold enrichment = 1.30) (**Fig. 5b**).

**Figure 5.**
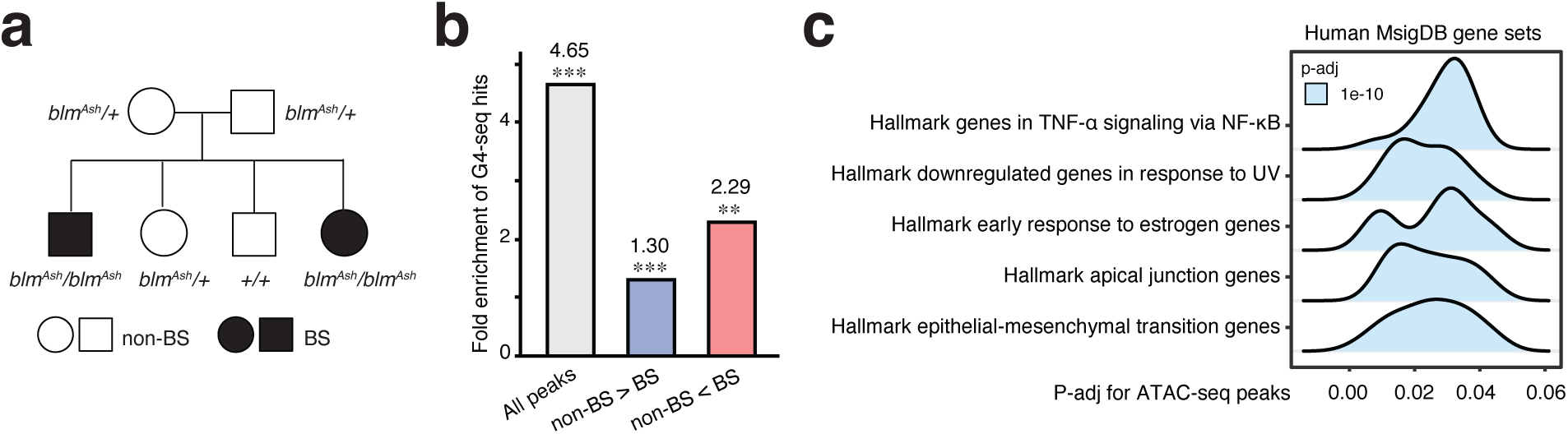
More accessible chromatin regions in Bloom Syndrome individuals were more enriched for G4-seq hits. **(a)** Pedigree of the Bloom Syndrome family. **(b)** Enrichment of G4-seq hits in differentially accessible chromatin regions. Z-test, ***, *P* < 0.001; **, *P* < 0.01. **(c)** Gene set enrichment analysis (hallmark pathways) of the differentially accessible chromatin regions.

We further analyzed the molecular functions of the differentially accessible chromatin regions. Notably, these peaks were enriched in genes related to TNF-α signaling via NF-κB, responses to UV, apical junction, and epithelial-mesenchymal transition (EMT) (**Fig. 5c**), potentially underlie the clinical symptoms. For instance, BS individuals are hypersensitive to UV light causing rashes on their faces ^1^. The EMT pathway plays an important role in morphogenesis and contributes to tumor invasion and metastasis. The dysregulation of this pathway may correspond to growth deficiency and increased risks of cancer in BS individuals. Notably, G-quadruplexes have been implicated in the regulation of EMT ^67^.

Regions with increased chromatin accessibility in BS individuals tend to have more G4-seq hits, echoing the extrapolated molecular regulatory mechanism from WT and BS cell lines. The epigenetic changes identified in BS individuals are associated with molecular functions relevant to the clinical symptoms in BS. Data from a Bloom Syndrome family also suggest G4s are prevalent in the molecular changes in BS, further highlighting the significance of G4 in the molecular etiology of BS.

## Discussion

Previous studies hinted at the relevance of G4s in the molecular changes in Bloom Syndrome by demonstrating possible association of G4-forming sequences, transcriptomic changes, and SCEs in BS cell lines ^21,24,25^. However, these studies showed mixed results, and they relied solely on *in silico* predicted G4 motifs in the human genome or *in vitro* validated G4-seq hits in purified genomic DNA—both of which fall short of actual G4 occurrence *in vivo*. Here, we carried out G4 ChIP-seq to map genome-wide endogenous G4 sites alongside RNA-seq and ATAC-seq in WT and BS lymphoblastoid and fibroblast cell lines. To our knowledge, our study is the first to map the endogenous G4s in BS and directly investigate the relationship between the local presence and change of G4s and BS.

Using cell lines derived from BS individuals and healthy donors, our data revealed a positive correlation of molecular changes in BS across all three modalities: endogenous G4 formation, open chromatin and gene expression. Next, using the small molecule PDS, we showed that G4 stabilization recapitulates much of the molecular phenotypes in BS (Fig. 3b). We further systematically leveraged various molecular signatures to gain mechanistic insights into G4s regulatory functions. Our data showed that G4 formation in open chromatin regions could further reinforce accessibility and G4 formation and accessible chromatin may act synergistically in enhancing transcription. Lastly, we collected data from a BS family and observed that more accessible chromatin regions in BS individuals showed stronger enrichment for G4-seq hits, further supporting the previously underappreciated role of G4 in BS. Our results for the first time provide direct evidence that one of the BLM’s substrates, G4s, is a central factor in linking BLM deficiency to molecular changes and clinical manifestations in BS.

Additionally, using SCE events, the signature of BS, mapped at single-cell resolution by van Wietmarschen et al. in the same LCL-BS cell line, we observed a significant enrichment of endogenous G4 sites at SCE locations, as well as in open chromatin regions (**Suppl. Fig.9**; *P* < 0.001, permutation, n=1000, see **Methods**). This provided direct evidence for the model that in BS, unresolved G4s stall DNA polymerase and initiates subsequent DNA repair, eventually resulting in elevated SCE frequencies ^11^.

Based on our findings, we propose a novel mechanistic model in which G-quadruplexes mediate molecular changes in BS cells and act as a central factor in the molecular etiology of Bloom Syndrome (**Fig. 6**). Firstly, during DNA replication, G4 formation hinders the progression of the DNA replication machinery leading to polymerase stalling. With intact BLM, G4s are resolved to ensure the progression of DNA replication. In contrast, in BLM-deficient cells, unsolved G4s cause DNA damage, initiate the homologous repair (HR) pathway, and eventually lead to sister chromatid exchanges due to the lack of BLM to dissolve the double Holliday junction in the last steps of HR. Secondly, an open chromatin state can create a self-reinforcing feedback loop between G4 and sustained open chromatin. In normal cells, G4s in the open chromatin regions are unwound such that histones can bind to DNA, switching the chromatin to a closed state. Conversely, in BS, unresolved G4 prevents nucleosome reassembly, causing the region to remain accessible and resulting in wider and more accessible chromatin regions. This altered nucleosome positioning subsequently influences gene expression by blocking certain factors (e.g. transcription factors, DNA methylation enzymes) from recognizing and binding to focal DNA. Further, G4s may recruit G4-binding proteins ^28,35,41^ and lastly, G-quadruplexes are likely to form in the promoter of highly expressed genes. During transcription in BS, unresolved G4 formation in the promoter enhances gene expression, which is likely through increasing the focal chromatin accessibility. Notably, G4 on the non-transcribing strand boosts the gene expression to a larger extent compared to G4 forming on the transcribing strand. These G4-mediated changes eventually contribute to the clinical phenotypes in BS individuals. As G4 landscapes are cell-type specific, also shown in our results, this model could provide insights into multifaceted symptoms across different tissues in BS.

**Figure 6.**
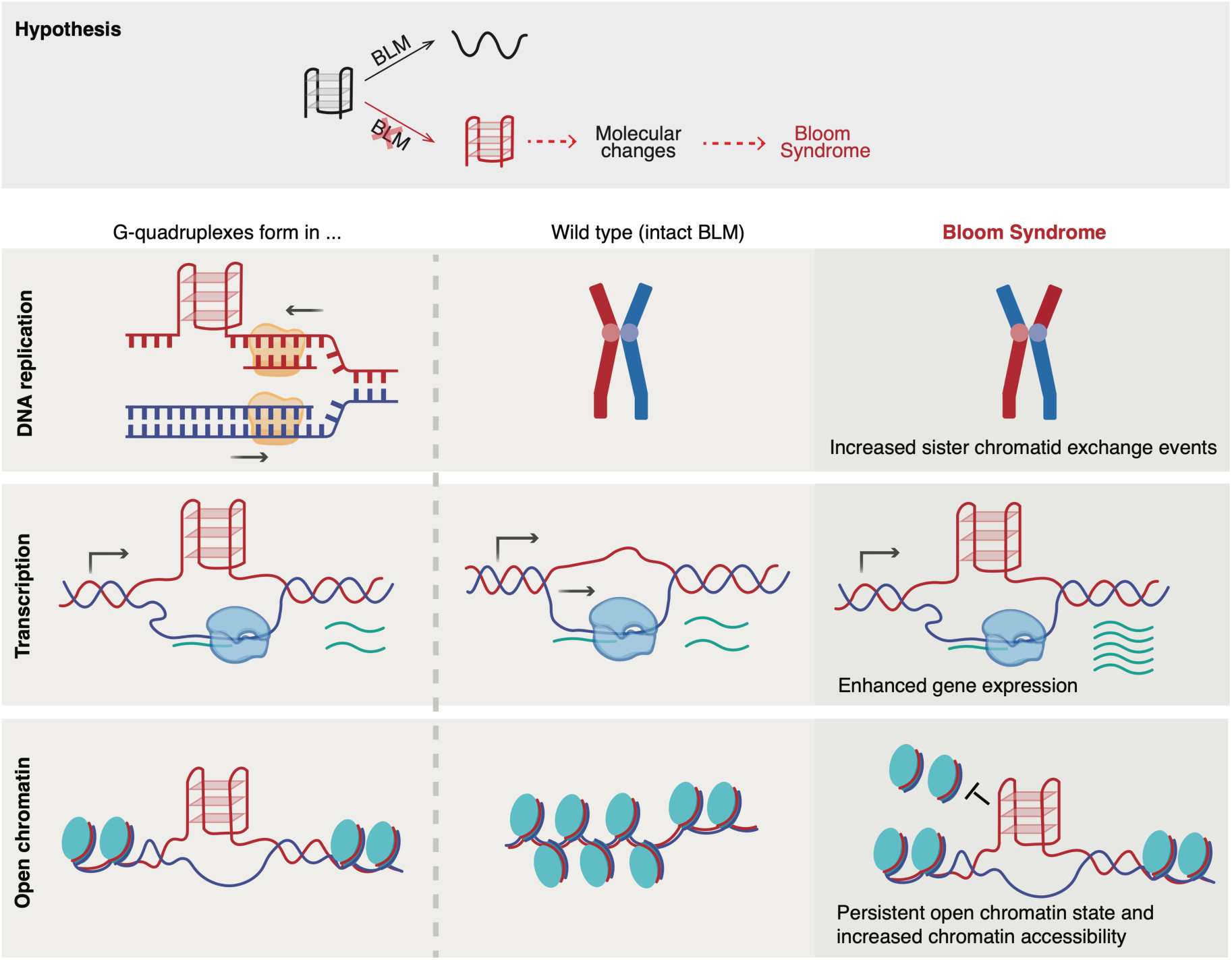
Mechanistic model of G4s in the molecular etiology of Bloom Syndrome. In Bloom Syndrome, defective G4-resolving activity leads to altered G4 formation, which in turn drives the molecular changes underlying the disease (hypothesis). During DNA replication, unresolved G4s stall replication forks, cause DNA damages, trigger DNA repair pathways, and ultimately result in elevated sister chromatid exchange events. During transcription, persistent G4s in gene promoters — particularly those forming on the non-transcribing strand—enhance gene expression. In parallel, unresolved G4s maintain chromatin in an open and accessible state, reinforcing transcriptional activation and further shaping gene regulatory outputs. In parallel, unresolved G4s maintain chromatin in an open and accessible state, reinforcing transcriptional activation and further shaping gene regulatory outputs.

Functional enrichment analyses on differential signals in BS cells, upon PDS treatment or in BS individuals further support the aforementioned mechanistic link between G4 formation and BS. Genes downregulated in BS cells or after PDS treatment were enriched in tRNA processing, non-coding RNA metabolism, and ribosome biogenesis—consistent with previous findings that BLM binds rDNA repeats and regulates rRNA transcription by RNA polymerase I via resolving G4s formed between DNA and RNA hybrids ^56,57,68^. In data both from cell lines and the BS family, differential signals showed functional enrichment for epithelial-mesenchymal transition (EMT). This process is characterized by the expression of certain cell adhesion molecules and essential for embryonic development and organogenesis ^69^. Notably, this process is reported to be regulated by G-quadruplexes ^67,70,71^. Dysregulation of EMT may underlie the growth deficiencies observed in BS and may also contribute to the increased cancer susceptibility, given EMT’s role in metastasis ^69^. Additionally, enrichment for UV response pathways aligns with BLM’s role in DNA repair and the clinical manifestation of UV hypersensitivity in BS patients ^1^.

Previous studies have reported conflicting results regarding the enrichment of G4-forming sequences, G4 motifs or G4-seq hits, at 1st-Ex-Int junctions of DE genes in BS ^24,25^. In our study, we directly addressed this question using G4 ChIP-seq peaks. We did not observe a consistent enrichment of G4s at the 1st-Ex-Int junction of the DE genes across both cell types. Since G4s are particularly enriched around TSSs, we further examined TSSs of DE genes and observed enrichment in only one cell type. Similarly, while background G4 enrichment at TSSs was observed in both cell types, enrichment at the TSSs of DE genes was detected only in fibroblasts and not in LCLs. (permutation, n = 1000; fibroblasts, *P* < 0.001; LCL, *P* = 0.20) (**Suppl. Fig. 3e-h**; see **Methods**). This reinforces the point that putative G4-forming sequences do not necessarily reflect *in vivo* G4 formation. The enrichment of G4s at TSS or 1st-Ex-Int junction of DE genes seems to be tissue-specific, underscoring the importance of mapping endogenous G4s within the appropriate cellular context.

A notable feature of endogenous G4s (here we mainly refer to the inter-molecular G4s. Intra-molecular G4s cannot be detected by G4 ChIP-seq and are not included in our discussion) is their strand-specificity. A previous *in vitro* study showed that G4 formation on the non-transcribing strand enhances transcription by promoting R-loop formation and elongation whereas G4 formation on the transcribing strand represses and even abolishes transcription ^33^. Although the G4 ChIP-seq assay does not capture strand-specific information, we inferred the strandedness of G4s by intersecting G4 peaks with strand-resolved G4-seq data. We showed that G4s are associated with enhanced gene expression regardless of strandedness, especially when G4s form on the non-transcribing strand while G4 strandedness had little impact on chromatin accessibility (**Suppl. Fig. 6f**). Our observation does not contradict the earlier *in vitro* studies because G4 ChIP-seq is a bulk assay and the detected G4 peaks only represent the G4 formation status in a proportion of the cells. Though in these cells the gene expression is largely reduced compared to the genes without G4 formation, the averaged signal from the bulk would remain higher than those genes without G4 peaks, though it is lower than the genes with G4 on the non-transcribing strand. It would be advantageous to detect G4 at single-cell levels to directly address how the strandedness of G4 affects gene expression.

Notably, G4 stabilization via PDS captured a substantial proportion (approximately 10% to 20%) of the changes observed in BS, despite several expected differences between PDS treatment and the disease context. Firstly, PDS treatment is short-term perturbation, whereas BS reflects the cumulative effects of long-term BLM deficiency. Secondly, G4 is one of BLM’s substrates, and G4 stabilization via PDS mimics the universal impairment of G4-resolving abilities, not specifically limited to those resolved by BLM if there are helicase-specific subsets of G4. Moreover, G4 dynamics are likely distinct between BS and PDS treatment. PDS strongly shifts the equilibrium of G4 toward G4 formation, and persistent G4 formation may render the focal chromatin rigid and impose even higher stress on molecular processes. While our results cannot fully address the causal role of G4, we could address that the relationship between G4 and molecular changes in BS is more than an association, underscoring the functional relevance of altered G4 dynamics in the BS. To fully establish the causal role of G4s, further studies and more sensitive assays for mapping G4s, such as CUT&Tag or even single-cell assays are required ^72,73^.

BLM helicase is not the only G4-unwinding enzyme, other helicases such as WRN, FANCJ, and DHX36 can also unwind DNA G-quadruplexes ^74,75^. We acknowledge that there might be partial redundancy between them. However, other helicases cannot fully compensate for BLM ^4^. Crucially, distinguished from others, BLM has a unique role in homologous recombination repair to suppress DNA recombination ^76^, and we observed that the excessive SCE events due to BS were enriched at G4 sites, further signifying the link between perturbed G4 dynamics and molecular phenotypes of BS. It would also be worth exploring the broader implications of G-quadruplexes in diseases caused by loss-of-function of other G4-unwinding helicases such as Werner Syndrome ^8^, to gain a deeper insight into the redundancy and specificity of different helicases for certain subsets of G4 structures.

In conclusion, we provided direct evidence that G4s could emerge as a central regulatory factor coordinating molecular phenotypes in BS. Based on the observed association, partial causation, and validation from BS individuals, we propose a molecular model whereupon the loss of function of BLM, defective G4-resolving abilities lead to G4 formation changes, resulting in subsequent molecular alterations in BS that contribute to the clinical phenotypes. Therefore, G4s may serve as a potential therapeutic target in BS. More broadly, this study enriches the understanding of Bloom Syndrome at a molecular level and the regulatory function of G4s *in vivo*. Additionally, we revealed a more complex molecular function of BLM that it possesses broad regulatory potential by regulating G4s, extending its functional impact beyond the canonical helicase activities.

## Methods

### Cell culture

Sex-matched and roughly age-matched fibroblast cell lines from Bloom syndrome and healthy donors were obtained from Coriell Institute. Lymphoblastoid cell lines (LCL) are GM16375 (*BLM* p.Ser595Ter) and AG14980 (*BLM*+/+); fibroblast cell lines are GM08505 (blm^Ash^, *BLM* c.2207_2212delinsTAGATTC) and GM00637(*BLM*+/+). LCLs were cultured in RPMI 1640 medium (Gibco, Cat. #21875) supplemented with 15% fetal bovine serum (FBS, Gibco, UK, Cat. #10500), 1% Penicillin-Streptomycin (Thermo Scientific, Gibco, Cat. #15140122) and 1% GlutaMAX-I (Gibco, UK, Cat. #35050). fibroblast lines were cultured in 1× DMEM (Thermo Scientific, Gibco, Cat. #11960) supplemented with 1% minimum essential medium non-essential amino acids (MEM NEAA; Thermo Scientific, Gibco, Cat. #11140) and 1% P/S and 10% or 15% fetal bovine serum (FBS; Thermo Scientific, Gibco, Cat. #10500064), respectively.

To collect cells for different experiments, fibroblast cell lines were first trypsinized (Trypsin-EDTA solution, Sigma-Aldrich, Cat. #T4049), pelleted at 300 g for 5 min, and washed once with cold 1x Dulbecco’s phosphate-buffered saline (PBS, Thermo Scientific, Gibco, Cat. #14190). Lymphoblastoid cells were pelleted at 300 g for 5 min and washed once with cold 1x PBS.

### Small molecule treatment

Pyridostatin (PDS, Sigma-Aldrich, Cat. #SML2690) stocks (2.5 mM) were prepared by dissolving it in DMSO (Sigma-Aldrich, Cat. #D2650). For LCLs, wild-type cells from cell line AG14980 were treated with 10 μM PDS vs. 0.4% DMSO (control) for 24 h at the concentration of 1 million cells per ml. Cells were then cultured with gentle rotation at 100 rpm to ensure equal exposure of cells to the treatment. For fibroblasts, wild-type cells (GM00637) at 60% confluency were treated with 10 μM PDS vs. 0.4% DMSO (referred as control) for 24 h.

### ATAC-seq library preparation

#### Cell lines

Cells were collected as described above and counted (automated cell counter Countesss 3, Thermo Scientific; Countess™ Cell Counting Chamber Slides Cat. # C10283). For each reaction 50, 000 cells were subjected to ATAC-seq preparation essentially as described in Corces, M. et al. ^77^ with modified tagmentation reaction. Nuclei were resuspended in 50 μl of tagmentation mix consisting of 10 μl 5× TAPS-DMF buffer (50 mM TAPS, 25 mM MgCl_2_, 50% (v/v) DMF), 3 μl Tn5 transposase ^78^, 16.5 μl 1x PBS, 0.25 μl 2% Digitonin, 0.5 μl 10% (v/v) Tween 20 and 19.75 μl H_2_O. DNA purified from the transposed product was then amplified via PCR (10 cycles) using Q5 High-Fidelity DNA Polymerase (New England Biolabs, Cat. #M0491L) and indexed primers N5xx and N7xx (synthesized by Integrated DNA Technologies) (Supplementary Table).

#### BS family samples

The procedure is the same as described above for lymphoblastoid exception for a longer cell lysis incubation on ice (10 min) and pelleting nuclei at 1000 g for 10 min.

### RNA extraction and RNA-seq library preparation

#### RNA extraction

Cell collection was performed as described above. The collected cells were lysed in 1 ml of Trizol (Sigma, Cat. #15596026) at room temperature (RT) for 5 min. Subsequently, 200 μl of chloroform (Thermo, Cat. #J67241.AP) was added, and the samples were centrifuged at 12,000 g for 15 min at 4°C to separate the phases. Prior to centrifugation, the samples were vortexed for 15 seconds and incubated at RT for 3 min. The upper phase, containing RNA, was carefully transferred to a new tube and mixed with 500 μl of isopropanol. The RNA was then precipitated at -20°C for 2 hours and subsequently pelleted at 12,000 g for 30 min at 4°C. The supernatant was discarded, and the RNA pellet was washed with 1 ml of ice-cold 75% ethanol. After centrifugation at 9,000 g for 5 min at 4°C, the ethanol wash step was repeated, and residual ethanol was aspirated using a pipette. The RNA pellet was air-dried by leaving the tubes open on the counter for approximately 5 min.The air-dried RNA pellet was resuspended in 50 μl of DNase reaction mix (5 μl of 10× Reaction Buffer for Turbo DNase, 1 μl of TURBO™ DNase (Invitrogen, Cat. #AM2238), 0.5 μl of recombinant RNase inhibitors (Jena Bioscience, Cat. #PCR-392S), and 43.5 μl of RNase-free water). The samples were incubated at 37°C for 25 min with gentle agitation at 550 rpm to carry out DNase treatment. For reextracting RNA, 150 μl of H_2_O and 200 μl of Phenol/Chloroform/Isoamylalcohol were added. The samples were vortexed for 15 s and then centrifuged at 13,000 rpm for 2 min at 4°C. The upper phase was transferred to a new tube and mixed with 20 μl of 3M Na-acetate pH 5.5 and 200 μl of isopropanol. The samples were incubated at -20°C for 1 hour to allow for the second RNA precipitation. The RNA was then pelleted, and the washing procedure was repeated as described in the initial precipitation step. The RNA pellet obtained after the final precipitation steps was dissolved in H2O.

#### RNA Quality assessment and library preparation

The quality of the RNA samples was assessed using Bioanalyzer RNA 6000 Nano assay (Agilent, Cat. # 5067-1511). Only RNA samples with a RIN (RNA Integrity Number) score exceeding 7 were selected for subsequent steps. RNA-seq library preparation was performed on 1 μg RNA using a poly-A mRNA enrichment kit (New England Biolabs, Cat. #E7490) and RNA-seq library preparation kit (New England Biolabs, Cat. #E7760S), following manufacturer’s recommendations.

### G-quadruplex ChIP-seq protocol and library preparation

#### Chromatin preparation for fibroblasts and LCL

Chromatin preparation was performed essentially as previously described with a few modifications ^47^. About 30 Mil ∼ 35 Mil LCL cells were collected and fixed with 0.5% formaldehyde (Thermo Scientific, Cat. #28906) in cell culture medium at RT for 6 min with gentle shaking. Cells were resuspended and treated in 1.5ml hypotonic buffer (Chromatrap, Cat. #100008) for 10 min. Nuclei were then lysed in 600μl nuclei lysis buffer (Chromatrap, Cat. #100008) and sheared in a 2ml sonication tube (Tube & Cap 12×24mm, Cat. # 520056) on a Covaries machine (Covaris S220) with the following parameters: *water level=15, duty cycle=18, intensity =7, cycle per burst = 200*. Each chromatin sample underwent shearing for a total of 15 min, with a 1 min pause for every 5 min of shearing.

For fibroblasts, cells from two full 15cm dishes were initially rinsed once in 1× PBS and subsequently fixed in the dish with 1% formaldehyde in cell culture medium at room temperature for 8.5 min. Nuclei were then lysed in 25 μl lysis buffer and sheared in small tubes (microTUBE AFA Fiber Pre-Slit Snap-cap 6×16mm, Covaris, Cat. #520045) using an S220 instrument (Covaris). The shearing parameters were as follows: *water level = 15*, *duty cycle = 15*, *intensity = 6*, and *cycle per burst = 200*. Each chromatin sample underwent shearing for a total of 3 min, with a 30-second pause for every one min of shearing.

#### G-quadruplex chromatin immunoprecipitation

For each sample, 2 biological replicates were prepared, each in turn divided into 3 technical replicates of immune-precipitation, except for the wild-type fibroblasts, for which the second biological replicate was performed with only 2 technical replicates due to the limited amount of BG4 antibody. The IP reaction was performed essentially as described previously ^47^, with the following modifications. During IP, chromatin was incubated with the BG4 antibody (Absolute antibody, Cat. #Ab00174-30.146) for an extended duration of 2 hours and double the amount of anti-FLAG M2 magnetic beads (Sigma-Aldrich, Cat. #M8823; equivalent to 10 μl of original beads) were used in the pull-down. To purify DNA from both the IP samples (DNA pulled down by the beads) and the input samples (sheared chromatin without going through IP steps) after 5 times of washing the beads with WASH buffer (100 mM KCl, 0.1% (v/v) Tween 20, 10 mM Tris, pH 7.4) to remove residual nonspecific bound chromatin, 75 μl of reverse-crosslinking buffer (0.2% SDS, 1× TE, and 50 mM NaCl) was introduced. The samples were incubated at 37°C for one hour, followed by an overnight incubation at 65°C. After an additional hour of proteinase K digestion at 65°C, DNA purification was carried out using a MinElute kit (QIAGEN, Cat. #28006). The subsequent steps for library preparation were carried out with the DNA Thruplex kit (Takara, Cat. #R400674, Cat. #R400665) following the manufacturer’s protocols.

#### G-quadruplex ChIP-qPCR and library preparation

Prior to library preparation, we used 2× CFX SYBR Mix (Applied Biosystems, Cat. #4472942) and a set of primers targeting known G4 positive and negative control sites to perform qPCR to evaluate G-quadruplex enrichment from IP vs. Input. Only samples with at least 5-fold enrichment of G4 positive sites (KLF14, GAPDH) to G4 negative sites (TMCC1) (**Supple. Table 5**) ^47^.

### Sequencing and data analysis

One biological replicate of G4 ChIP-seq libraries from GM08505 and GM00637 were sequenced on a Hiseq3000 platform (paired-end sequencing of 150 nt each; Illumina Inc.; service provider: Genome center in the Max Planck Institute for Biology, Tübingen). The rest of the samples were sequenced on a NovaSeq6000 platform (paired-end sequencing of 150 nt each; Illumina Inc., San Diego, etc.; service provider: GENEWIZ, Leipzig)

### ATAC-seq data analysis

#### Mapping and peak calling for ATAC-seq

Raw FASTQ reads were extracted with *bcl2fastq* (version 2.20). Tn5 and Truseq adapter sequences, G repeats due to sequencing artefacts, and reads shorter than 20 bp were trimmed and removed using *cutadapt* (version 4.0) ^79^. Reads were then aligned to human reference genome version hg38 or composite genome of hg38 and EBV using *bwa mem* (Version: 0.7.17-r1188) ^80^. Duplicates were marked and removed using *Picard* (version 2.18.25) (“Picard Toolkit.” 2019. Broad Institute, GitHub Repository. https://broadinstitute.github.io/picard/). Prior to calling peaks, mitochondrial reads, reads with a mapping quality lower than 20, and reads in hg38 blacklisted regions (http://mitra.stanford.edu/kundaje/akundaje/release/blacklists/hg38human/hg38.blacklist.bed.gz) were discarded.

ATAC-seq peak calling was performed using *MACS2* (version 2.1.1.20160309) with *callpeak –format BAMPE --nomodel –min-length 100 narrowPeak* parameters ^58^. For a given cell type, two types of ATAC-seq peak sets were generated. Using LCL as an example, to identify differentially accessible regions in each contrast (BS vs. WT or PDS vs. control), a local peak set was called by pooling an equal number reads from each sample. To identify peaks that are differentially accessible in BS and PDS–treated conditions, equal number of reads from WT, BS, control and PDS–treated samples from the same cell type was pooled to call a global peak set, which was further supplemented by unique peaks in the local peak sets from BS vs. WT or PDS vs. DMSO contrasts.

#### Differential analysis

To identify differential signals in BS vs. WT, the number of fragments in each ATAC-seq peak was counted with *htseq-count* (HTSeq version 0.9.1) ^81^. Subsequently, the count matrix was normalized and differential analysis was carried out in *DESeq2* (version 1.30.1) to identify significantly differential signals (adjusted p-value < 0.05) ^54,82^. In the case of applying copy number normalization, the count matrix was adjusted before data normalization as described in the following section. However, we noticed that the normalization method in *DESeq2* produced a strongly skewed MA plot when comparing PDS–treated and control LCL samples. We thus employed the loess normalization method in *csaw* (version 1.24.3) following differential analysis with *edgeR* (version 3.32.1) for comparing PDS– vs. DMSO-treated samples ^83,84^, as the method IV described in Yan et al. ^85^.

#### Copy number normalization

Copy number normalization is done as described in Su et al. ^55^. Briefly, prior to the differential analysis of ATAC-seq, copy number normalization was applied to the read count for peaks. The first step of copy number normalization is characterizing CNR. Genomic sequencing data from cell lines GM08505 (BS) and GM00637 (WT) were used in *CNVkit* following its recommended copy number calling pipeline *(--method wgs --target-avg-size 50000*) to identify copy number alteration in BS relative to WT ^86^. Gapped regions in the human genome assembly marked by Ns (http://hgdownload.soe.ucsc.edu/goldenPath/hg38/database/gap.txt.gz) were excluded from the analysis. In the output from *CNVkit*, log_2_-transformed values representing copy number ratio (CNR) for genomic segments were retrieved and converted back to the original value. The second step is assigning each peak to its overlapping DNA segment or the closest DNA segment. The CNR of this segment was then used as a scaling factor to modify the read/fragment count in this peak. Specifically, for the BS vs. WT comparison, if the CNR is greater than or equal to 1 (CNR ≥ 1), the fragment counts in peaks in BS were divided by the CNR. Conversely, if the CNR was smaller than 1 (CNR < 1), the fragment counts in peaks in WT were multiplied by the CNR.

### RNA-seq data analysis

#### Differential analysis of gene expression

RNA-seq reads from fibroblast samples were aligned to hg38 with gtf file (comprehensive gene annotation file for regions CHR from release 40 (GRCh38.p13, https://www.gencodegenes.org/human/release_40.html) and STAR (version 2.7.9a) with default parameters ^87^. RNA-seq reads from LCL samples were aligned to a composite reference genome of human and EBV (EBV genome is downloaded from NCBI, NC_007605.1) with the corresponding composite gtf files (gff3 for EBV is created from NC_007605.1) via *STAR* ^87^. Taking advantage of the directionality of the RNA-seq library preparation protocol, the count of reads mapping to the reverse strand was used for subsequent differential analysis in *DESeq2* with default parameters. Transcripts per million (TPM) is used as a measure of normalized gene expression values and genes were defined as expressed if they have z-standardized TPM values greater than -3 ^88,89^.

### ChIP-seq data analysis

#### Peak calling and filtering

We processed the reads until the step of read filtering in the same way as described in the ATAC-seq reads processing for LCL and fibroblasts. For each biological replicate, a pooled file was generated by subsampling 35 million reads from each technical replicate. The pooled file was subjected to peak calling using *MACS2* (version 2.1.1.20160309) with *callpeak –format BAM narrowPeak* ^58^. Peak sets from two biological replicates were ranked by their signal values and then filtered based on reproducibility using *IDR* (version 2.0.3) ^59^. The final high confidence peaks for each sample were obtained with the criteria of irreproducible discovery rate (IDR) < 0.05. Peaks were also called with input samples using the same parameters, which were later used as regions to be excluded in the differential analysis.

#### Differential analysis

The differential analysis was carried out in *DiffBind* ^56^. Signals were quantified with default parameters *summits = FALSE, bFullLibSize = FALSE* followed by data normalization and statistical test in *edgeR*. We define significantly differential G4 forming sites as those with FDR < 0.05. For fibroblast-BS vs. fibroblast-WT, after signal quantification and before data normalization, the read count was extracted from *DiffBind* and subjected to copy number normalization, as described in Su et al. ^63^.

### Peak annotation and functional enrichment analysis

ATAC-seq and ChIP-seq peaks were annotated in *ChIPseeker* with the gtf file used in RNA-seq read alignment with default parameters ^90^. Gene Ontology enrichment analysis on gene lists was carried out with *ChIPseeker* and *Reactome* in *R* ^90,91^. Alternatively, significantly DE genes were ranked by the fold change and subjected to gene set enrichment analysis against the human hallmark gene sets with *msigdbr* in *R*.

### Peak signal profiles

Bamfiles were converted to bigwig format via *bamCoverage* in *deeptools* with the option *--binSize 20 -of bigwig --normalizeUsing RPKM* ^92^. Subsequently, bigwig files were used to compute the signal in regions of interest (TSS±3kb, gene body) via *computeMatrix* function followed by function *plotHeatmap* and/or *plotProfiles* in *deeptools* with default parameters.

### Principal component analysis (PCA)

A read count matrix with the raw count was created by summarizing the fragment count in peaks for ATAC-seq and ChIP-seq and the read count per gene for RNA-seq. The read count matrix was then transformed with the *DESeq2* function *varianceStabilizingTransformation*, following the steps outlined in the *DESeq2* manual ^54^. Subsequently, PCA analysis was carried out using the *R* function *prcomp* with the option *scale = FALSE* on the processed read count matrix. The coordinate and the percentage of variance explained by each component were extracted and plotting was done with *ggplot2*.

### Converting ATAC-seq, RNA-seq and ChIP-seq signals to z-scores

For ATAC-seq and ChIP-seq, we first converted raw fragment count in peaks to RPKM (reads per kilobase per million reads) using the total number of fragments in peaks as the total library size. For RNA-seq, we converted the fragment count per gene to TPM (transcripts per million) using the mean transcript length of each gene ^88^. RPKM and TPM values are then converted to z score via *zFPKM* package in R with the function *zFPKM* ^89^. Genes whose expression level has a z-score of smaller than -3 is defined as inactive or not expressed ^89^.

### Calcualting the density of ATAC-seq peaks

We counted the number of ATAC-seq peaks in the flanking 10kb region of each ATAC-seq peak. Then we stratified ATAC-seq peaks into those overlapping G4 peaks (G4+ ATAC-seq peaks) and those not overlapping G4 peaks (G4− ATAC-seq peaks) and compared their peak densities.

### Correlation between differential signals in RNA-seq and ATAC-seq or ChIP-seq

In this study, the promoter is defined as the upstream 3kb (-3kb) to downstream 3kb (+3kb) region of a transcription start site (TSS). A gene will be assigned as the target gene of an ATAC-seq peak or G4 peak if the peak is less than 3kb away from the TSS. Correlation between them is assessed by comparing log_2_FC values of RNA-seq and ChIP-seq for genes with G4 peak in their promoter in *cor.test*.

### Correlation between differential signals in ATAC-seq or ChIP-seq

ATAC-seq and ChIP-seq peaks were intersected. An ATAC-seq peak is considered as a potential target of a G4 ChIP-seq peak if their overlap is larger than 150bp. Pearson correlation between them is tested with log_2_FC values of ATAC-seq and ChIP-seq in *cor.test*.

### Inferring the strandedness of G4 peaks

We downloaded the G4-seq hits identified in purified human genomic DNA upon the addition of K^+^ in the sequencing buffers via G-quadruplex induced polymerase stalling (GEO: GSE63874, https://www.ncbi.nlm.nih.gov/geo/query/acc.cgi?acc=GSE63874, GSE63874_Na_K_minus_hits_intersect.bed.gz, GSE63874_Na_K_plus_hits_intersect.bed.gz) and lifted their coordinate to hg38 via *LiftOver* with the corresponding chain file (https://hgdownload.cse.ucsc.edu/goldenpath/hg19/liftOver/hg19ToHg38.over.chain.gz) ^43,93^. A G4 peak is postulated to form on the Watson strand and if the G4 peak only intersects G4-seq hits on Watson strand. The same principle applies to identifying G4s on Crick strand. Based on the strandedness of the gene that the G4 peaks is assigned to, it can be inferred whether the G4 structure forms on the transcribing or non-transcribing strand.

### Testing interaction between G4 peaks and ATAC-seq

We tested the relationship between differential gene expression(diff-RNA), differential chromatin accessibility (diff-ATAC), and differential G4 formation (diff-G4). We used the *Car* package in *R* and built a multifactor ANOVA model with interactions and the *type-III* test. The model was *diff-RNA = diff-ATAC + diff-G4 + diff-ATAC x diff-G4*. Values for diff-RNA is the actual log_2_FC whereas diff-ATAC and diff-G4 are categorical values of their differential status, non-differential, up-regulated (p-adj < 0.05 or FDR < 0.05 and log_2_FC(BS/WT) > 0), or down-regulated (up-regulated (p-adj < 0.05 or FDR < 0.05 and log_2_FC(BS/WT) < 0). We then carried *type-III* tests, a traditional multifactor ANOVA model with interactions.

### Enrichment of G4 peaks at certain genetic features

Enrichment is assessed by comparing the observed overlap between G4 peaks and certain genetic features to the expected number of overlaps by chance. Using TSS as an example, to evaluate if the G4 peaks are enriched at the TSS of expressed genes, we first calculated the overlap between G4 peaks and TSS of expressed genes. Then we randomly permuted the genomic locations of G4 peaks in the genome with *bedtools shuffle* and counted the number of overlaps between randomly shuffled G4 peaks and TSS. The expected number of overlaps is represented by the mean number of overlaps in the permutation for 1000 times. We evaluated if the G4 peaks are particularly enriched at the TSS of DE genes via the function *permTest()* in *regionR* with the following parameter, *randomize.function=resampleRegions, evaluate.function=numOverlaps.* The TSS of DE genes were permuted in the TSS set of all expressed genes for 1000 times.

### Summary of statistical tests

All statistical tests were carried out in *R*. Person correlation was tested between ATAC log_2_FC and RNA log_2_FC, between G4 ChIP log_2_FC and ATAC log_2_FC, and between G4 ChIP log_2_FC and RNA log_2_FC (Fig. 2a, 2b, 2d; Fig 4g; Suppl. Fig. 1b,1d; Suppl. Fig. 4a, 4b, 4d; Suppl. Fig. 6e,6g; Suppl. Fig. 7c, 7d).

Wilcoxon tests were used to compare the open chromatin accessibility, size and density of ATAC-seq peaks partitioned by G4 presence, gene expression levels partitioned by G4 presence in the promoter, gene expression levels stratified by the strandedness of G4, effect size of chromatin accessibility and G4 formation changes on gene expression changes (Fig. 4a, 4b, 4d, 4e, 4f; Fig 5h; Suppl. Fig. 6f; Suppl. Fig. 7b).

Z-tests were carried out in assessing if G4 peaks or G4-seq hits are enriched in certain genomic intervals or genetic features by comparing observed values and expected values (Fig. 1f; Fig. 5b; Suppl. Fig. 2g; Suppl. Fig. 3; Suppl. Fig.9). The fold enrichment is the ratio of observed values to expectested values. Expected values were estimated by permutation (n=1000).

Hypergeometric tests (*phyper* in *R*) were performed to test if there is an enrichment of BS+/PDS+ and BS–/PDS– signals with *lower.tail = FALSE* and if there is a depletion of BS+/PDS– and BS–/PDS+ signals with *lower.tail = TRUE* (Fig. 3c, 3d).

## Supplementary information

Supplementary figures 1-9

Supplementary tables 1-4

## Availability of data and materials

ATAC-seq and G4 ChIP-seq data from WT and BS fibroblast cell lines are available at the NCBI GEO repository under accession numbers GSE259257 and GSE259258.

ATAC-seq data from WT and BS fibroblast cell lines, WT fibroblast cell lines treated with DMSO or PDS and WT lymphoblastoid cell lines treated with DMSO or PDS are available at the NCBI GEO repository under accession numbers GSE283477.

ATAC-seq data from the Bloom Syndrome family members are available at the NCBI GEO repository under accession number GSE283480.

RNA-seq data in this study are available at the NCBI GEO repository under accession number GSE283485.

G4 ChIP-seq data from WT and BS lymphoblastoid cell lines are available at the NCBI GEO repository under accession numbers GSE283482.

Codes for the analyses are available at GitHub, https://github.com/Dingersrun/Copy-number-normalization and https://github.com/Dingersrun/G4-in-Bloom-Syndrome.

## Acknowledgments

We acknowledge the patients and families who provided their samples to the Bloom syndrome Registry, without whom this work would not be possible. Bloom syndrome Registry was supported, in part, by the New York Community Trust, United States, Weill Cornell Medicine’s Clinical and Translational Science Center, United States, and the National Center for Advancing Translational Science of the National Institutes of Health, United States Under Award Number UL1TR000457. We thank members in Frank Chan and Felicity Jones’ lab for helpful discussions. We thank Yinan Wang, Detlef Weigel, and Marja Timmermans for their input. We also thank the Genome Center at the Max Planck Institute for Biology Tübingen for providing support. We are grateful to Linda Chen for her support in graphic design. We thank Marek Kucka for bioinformatic support. We thank Andre Noll for computing support.

## Ethics declarations

### Ethics approval

Not applicable.

