## Supplementary Figures for "G-quadruplexes regulate chromatin accessibility and gene expression in Bloom Syndrome"

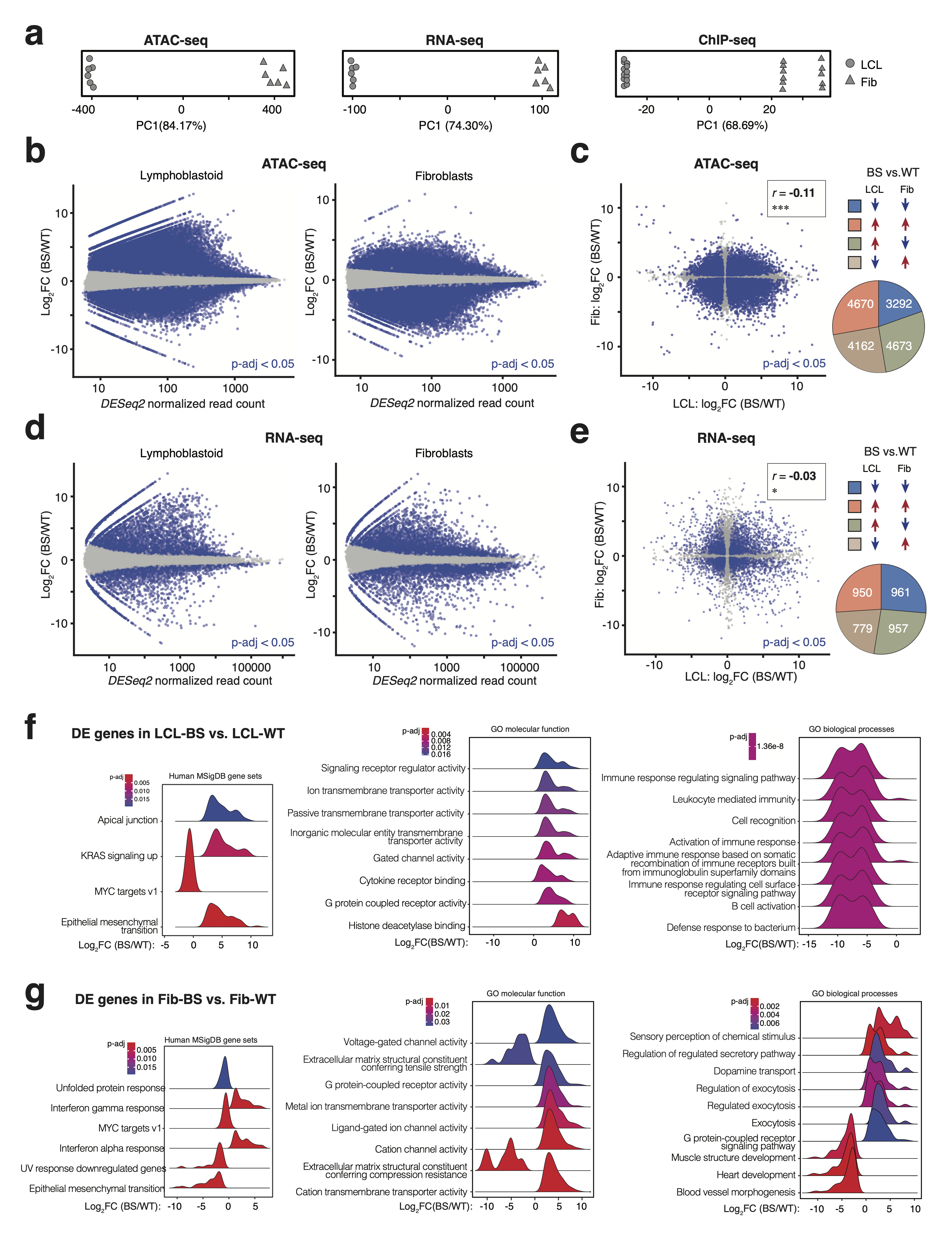
**Supplementary Figure 1. Changes in chromatin accessibility, gene expression and G4 formation in Bloom Syndrome cells.**

(**a**) Principal Component (PC) 1 of ATAC-seq, RNA-seq and G4 ChIP-seq data from WT and BS samples. LCL, lymphoblastoid cell lines; Fib, fibroblast cell lines.

(**b**) MA-plots of differentially accessible chromatin regions in LCL-BS vs. LCL-WT (left) and Fib-BS vs. Fib-WT (right). Peaks with p-adj < 0.05 are denoted in blue and those with p-adj ≥ 0.05 in gray.

(**c**) Comparison of differential ATAC-seq peaks in LCL-BS vs. LCL-WT and Fibroblast-BS vs. fibroblast-WT. Peaks with significant chromatin accessibility changes in both comparisons are denoted in blue and the rest in gray (left). Blue and red arrows indicate significant decreases and increases in the specified contrast, respectively (right). R is the Pearson correlation coefficient and ***, *P* < 2 × 10^-16^.

(**d**) MA-plots of differentially expressed genes in the RNA-seq analysis. Left, LCL-BS vs. LCL-WT; right, fibroblast-BS vs. fibroblast-WT. Significantly differentially expressed genes with p-adj < 0.05 are denoted in blue and those with p-adj ≥ 0.05 in gray.

(**e**) Comparison of differentially expressed genes in LCL-BS vs. LCL-WT and fibroblast-BS vs. fibroblast-WT. Significantly differentially expressed genes in both comparisons are denoted in blue and the rest in gray (left). Blue and red arrows indicate significant decreases and increases in the specified contrast, respectively (right). R is the Pearson correlation coefficient and *, *P* < 0.05.

Gene ontology and Gene set enrichment analysis with the significantly differentially expressed genes ranked by fold change in LCL-BS compared to LCL-WT (**f**) and fibroblast-BS compared to fibroblast-WT (**g**).

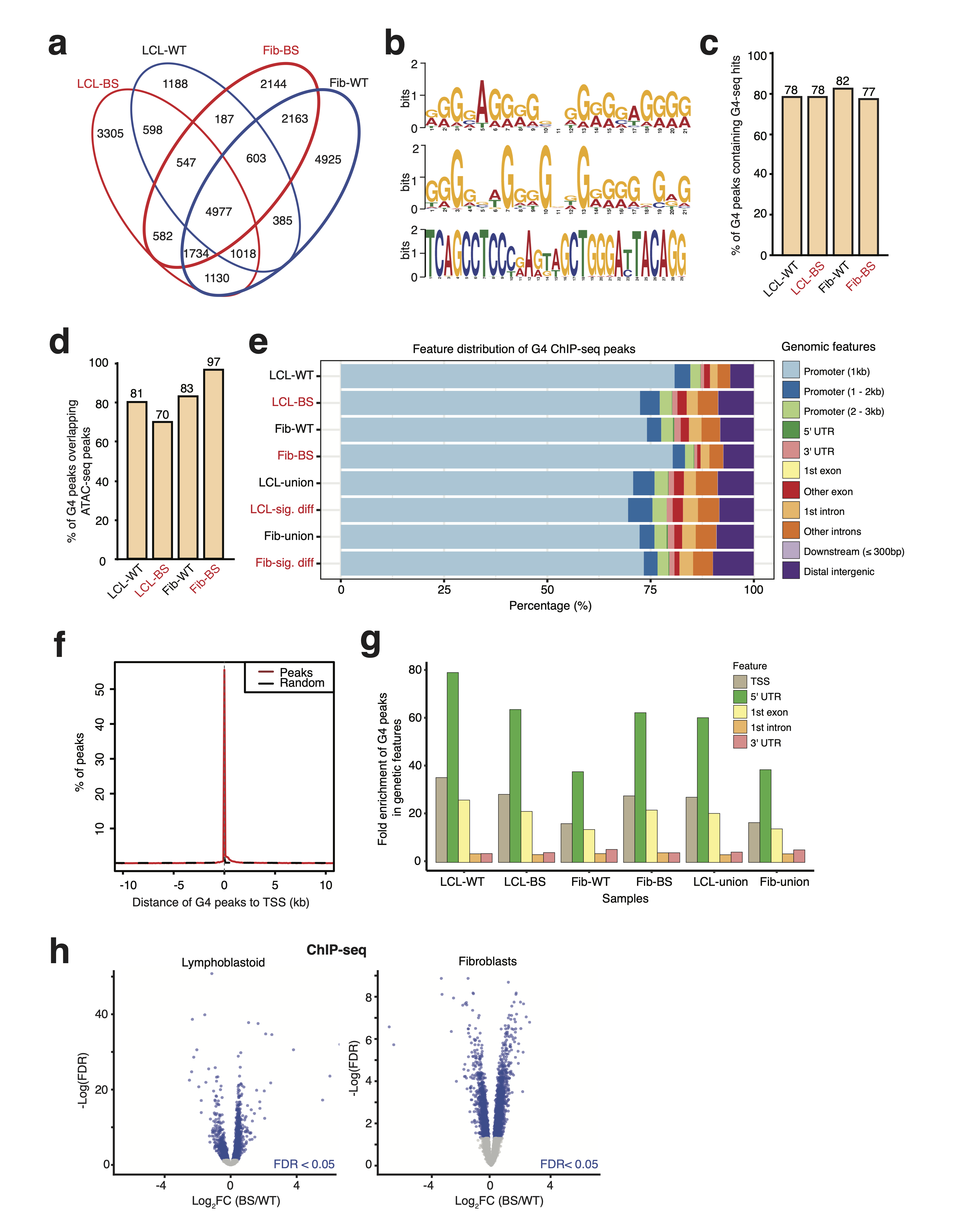
 **Supplementary Figure 2. Features of endogenous G4 peaks.**

(**a**) Number of G4 peaks in samples. LCL, lymphoblastoid cell lines; Fib, fibroblast cell lines.

(**b**) Examples of enriched DNA motifs detected by MEME in G4 peaks. For all 3 motifs, e-value < 1 × 10^-15^.

(**c**) Percentage of G4 peaks overlapping with G4-seq hits.

(**d**) Percentage of G4 peaks overlapping open chromatin regions.

(**e**) Distribution of G4 peaks in different genomic features. UTR, untranslated regions; union, the union of G4 peaks from both WT and BS cell lines; sig. diff, significantly differential G4 peaks in BS vs WT.

(**f**) Enrichment of G4 peaks at TSS.

(**g**) Fold enrichment of G4 peaks in different genomic features. Fold enrichment is the ratio of the observed overlap between G4 peaks and the genomic features to the expected overlap estimated by permutation (n=1000; z-test, for all features, *P* < 0.01; see **Methods**).

(**h**) Volcano plots of differentially G4-forming sites. Left, LCL-BS vs. LCL-WT; right, Fib-BS vs. Fib-WT. Peaks with FDR < 0.05 are denoted in blue and those with FDR ≥ 0.05 in gray.

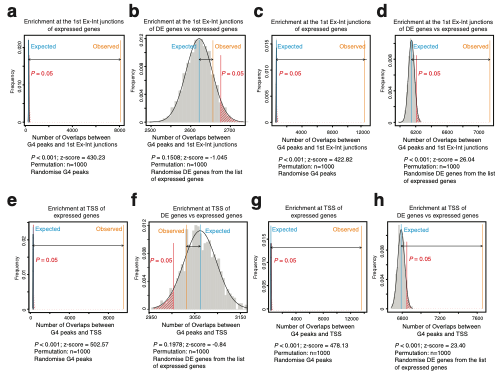
**Supplementary Figure 3. Enrichment of G4 peaks at the first exon-intron junctions and transcription start site of differentially expressed genes in Bloom Syndrome cells.**

Observed and expected number of G4 peaks intersecting the first exon-intron junctions (1st-Ex-Int) of expressed gene (**a**, LCL; **c**, fibroblasts) and of differentially expressed genes (**b**, LCL; **d**, fibroblasts). The number of observed and expected G4 peaks intersecting the transcription start site (TSS) of expressed gene (**e**, LCL; **g**, fibroblasts) and of differentially expressed genes (**f**, LCL; **h**, fibroblasts). Z-tests were carried out to compare the number of observed overlaps and expected overlaps by chance (permutation, n=1000; see **Methods**).

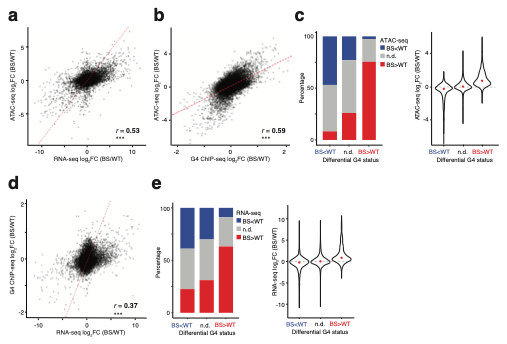
**Supplementary Figure 4. Differential G4 formation positively correlates with differential chromatin accessibility and differential gene expression in fibroblast Bloom Syndrome cells.**

For (**a**), (**b**) and (**d**), *r* is the Pearson correlation coefficient; ***, *P* < 2×10^-16^. (**a**) Positive correlation between differential chromatin accessibility and differential gene expression in fibroblast-BS vs. fibroblast-WT. The red line indicates the fitted linear regression line. Fib, fibroblast.

(**b**) Positive correlation between differential chromatin accessibility and differential G4 formation in LCL-BS vs. LCL-WT. The red line indicates the fitted linear regression line.

(**c**) Distribution of differentially accessible chromatin regions and their fold changes stratified by changes in G4 formation in fibroblast-BS vs. fibroblast-WT. Red dots denote the median log_2_FC.

(**d**) Positive correlation between differential gene expression and differential G4 formation in fibroblast-BS vs. fibroblast-WT. The red line indicates the fitted linear regression line.

(**e**) Distribution of differentially expressed genes and their fold changes stratified by changes in G4 formation in fibroblast-BS vs. fibroblast-WT. Red dots denote the median log_2_FC.

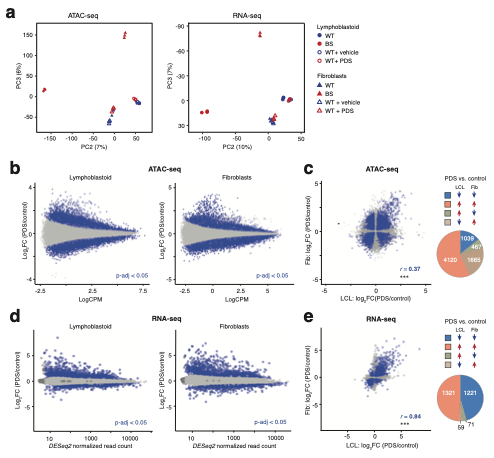
**Supplementary Figure 5. Changes in chromatin accessibility and gene expression in wild-type cells caused by PDS treatment.**

(**a**) Principal Component Analysis on the global open chromatin landscape (left) and the gene expression profile (right) of WT (filled blue symbols), BS (filled red symbols), WT treated with vehicle (unfilled blue symbols), and WT samples treated with PDS (unfilled red symbols).

(**b**) MA-plots of differentially accessible ATAC-seq peaks upon PDS treatment in LCL (left) and fibroblasts (Fib) (right). Significantly differentially expressed genes with p-adj < 0.05 are denoted in blue and those with p-adj ≥ 0.05 in gray.

(**c**) Comparison of differentially accessible ATAC-seq peaks upon PDS treatment in LCL and Fib. Significantly differential signals in both cells are denoted in blue and the rest in gray (left). Blue and red arrows indicate significant decreases and increases in the specified contrast, respectively (right). R is the Pearson correlation coefficient for the blue data points.

(**d**) MA-plots of differentially expressed genes upon PDS treatment in LCL (left) and fibroblast-(right). Significantly differentially expressed genes with p-adj < 0.05 are denoted in blue and those with p-adj ≥ 0.05 in gray.

(**e**) Comparison of differentially expressed genes upon PDS treatment in LCL and fibroblasts. Significantly differential signals in both cells are denoted in blue and the rest in gray (left). Blue and red arrows indicate significant decreases and increases in the specified contrast, respectively (right).

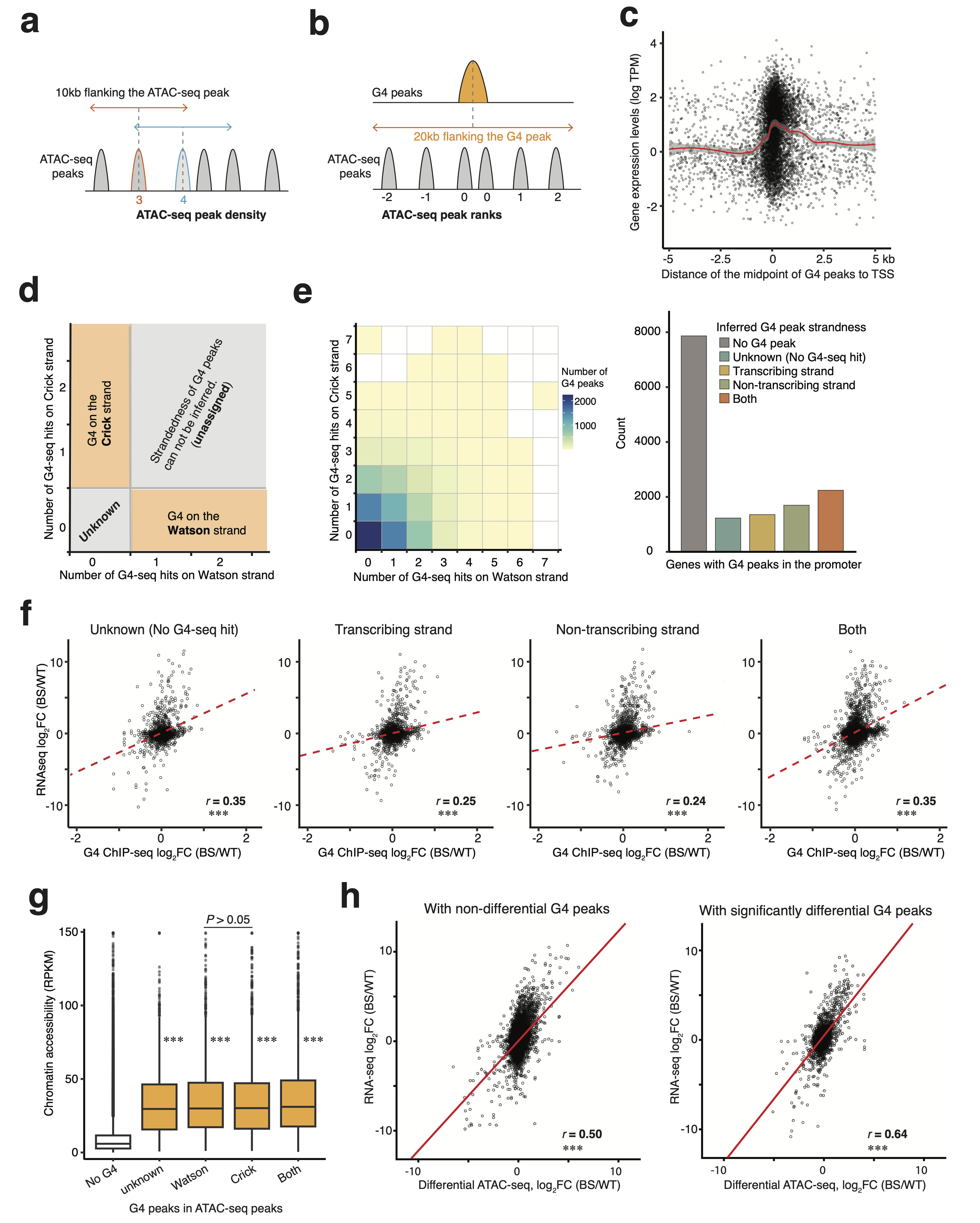
**Supplementary Figure 6. Impacts of G4 formation on chromatin accessibility and gene expression changes.**

(**a**) Schematic representation of estimating ATAC-seq peak density by counting the number of ATAC-seq peaks in 10 kb regions flanking the midpoint of an ATAC-seq peak.

(**b**) Schematic representation of ranking ATAC-seq peaks proximal to G4-peaks. Overlapping ATAC-seq peaks are ranked 0. Upstream and downstream ATAC-seq peaks have negative and positive ranks, respectively.

(**c**) The gene expression levels tending to decrease as the G4 peaks form further away from the TSS. The red line denotes the LOESS (locally estimated scatterplot smoothing) fitted curve for the data and the grey area depicts the 95% confidence interval for the fitted curve.

(**d**) Inferring the strandedness of G4s by intersecting G4 ChIP-seq peaks with strand-resolved G4-seq hits. G4 in G4 peaks containing G4-seq hits only from Watson (or Crick) strand were inferred to form on Watson (or Crick) strand. G4 peaks overlapping G4 seq on both strands are labeled as unassigned.

(**e**) Example of assigning the strandedness of G4 peaks, represented by the LCL-WT sample (left). Chromatin accessibility stratified by inferred strandedness of the G4s (right).

(**f**) ATAC-seq peaks overlapping G4 peaks show higher chromatin accessibility. Wilcoxon tests were used to compare each group with ATAC-seq peaks lacking G4 overlap (No-G4), unless otherwise specified; ***, *P* < 2 ×10^-16^.

(**g**) Correlation between G4 formation and gene expression changes stratified by the strandedness of the G4 relative to the transcribing strand. The red line indicates the fitted linear regression line. R is the Pearson correlation coefficient. ***, *P* < 2 ×10^-16^.

(**h**) Correlation between differential chromatin accessibility and differential gene expression stratified by the differential status of G4 peaks. The red line indicates the fitted linear regression line. R is the Pearson correlation coefficient. ***, *P* < 2×10^-16^.

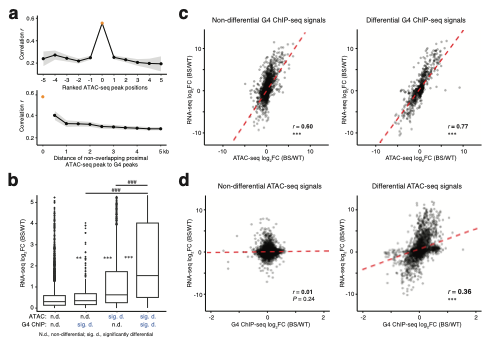
**Supplementary Figure 7. Molecular mechanisms of G4 regulating molecular changes in lymphoblastoid.**

(**a**) Correlation between G4 ChIP-seq log_2_FC and ATAC-seq log_2_FC of G4-overlapping or proximal ATAC-seq peaks.

(**b**) Additive effects of differential chromatin accessibility and G4 formation on differential gene expression in LCL-BS vs. LCL-WT. Wilcoxon tests were used. Groups with P values denoted by * were compared to the gene without G4 (the most left boxplot); **, *P* = 0.003, *P* < 2 × 10^-16^; ***, *P* < 2 × 10^-16^; ###, *P* < 2 × 10^-16^ .

(**c**) Correlation between differential gene expression and differential chromatin accessibility stratified by differential G4 status. The red line indicates the fitted linear regression. R is the Pearson correlation coefficient. ***, *P* < 2 × 10^-16^.

(**d**) Correlation between differential gene expression and differential G4 formations stratified by the differential status of chromatin accessibility. The red line indicates the fitted linear regression. R is the Pearson correlation coefficient. ***, *P* < 2 × 10^-16^.

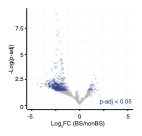

**Supplementary Figure 8. Differentially accessible chromatin regions in Bloom Syndrome individuals.** Significantly differentially expressed genes with p-adj < 0.05 are denoted in blue and those with p-adj ≥ 0.05 in gray.

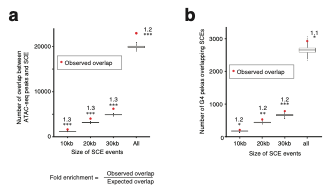

**Supplementary Figure 9. Enrichment of sister chromatid exchange (SCE) events at G4 sites and open chromatin regions.**

(**a**) Enrichment of sister chromatid exchange (SCE) events in LCL-BS at open chromatin regions in LCL-BS. Z-tests were carried out to compare observed overlap and expected overlap (permutation, n=1000; see **Methods**). ***, *P* < 0.001.

(**b**) Enrichment of sister chromatid exchange events at endogenous G4 forming sites in LCL-BS. Z-tests were carried out to compare observed overlap and expected overlap (permutation, n=1000; see **Methods**). *, *P* < 0.05; **, *P* < 0.01; ***, *P* < 0.001.
