## Supplementary Tables for "G-quadruplexes regulate chromatin accessibility and gene expression in Bloom Syndrome"

|  | Number of over lap with G4 peaks | | | | Number of overlap with ATAC-seq peaks | | | |
| --- | --- | --- | --- | --- | --- | --- | --- | --- |
|  | AllSCE | SCE_10kb | SCE_20kb | SCE_30kb | AllSCE | SCE_10kb | SCE_20kb | SCE_30kb |
| Observed | 2927 | 209 | 530 | 785 | 22934 | 1648 | 4042 | 6225 |
| Expected | 2644.5 ± 109.1 | 176.5 ± 14.8 | 438.5 ± 25.3 | 665.7 ± 33.1 | 19906.8 ± 320.0 | 1231.6 ± 38.2 | 3177.4 ± 70.5 | 4891.5 ± 88.5 |
| Fold enrichment | 1.11 | 1.18 | 1.21 | 1.18 | 1.15 | 1.34 | 1.27 | 1.27 |
| Z score | 2.55 | 2.20 | 3.62 | 3.61 | 9.73 | 10.91 | 12.27 | 15.06 |
| *P* | 1.40E-02 | 2.20E-02 | 2.00E-03 | 9.99E-04 | 9.99E-04 | 9.99E-04 | 9.99E-04 | 9.99E-04 |

**Supplementary Table 1. Enrichment of sister chromatin exchange events in G4 peaks and ATAC-seq peaks.**

Expected number of overlaps is estimated by permutation (n=1000). The number is represented as mean ± standard deviation

**Supplementary Table 2. Hypergeometric tests of enrichment and depletion of overlapping differentially accessible chromatin regions upon BLM deficiency and PDS treatment.**

| DA peak categories | Observed | Expected by chance | Fold enrichment | *P* (Hypergeometric test) |
| --- | --- | --- | --- | --- |
| LCL: BS- PDS- | 7608 | 3707 | **2.05** | 9.12E-2537 |
| LCL: BS+ PDS+ | 4446 | 4715 | 0.94 | 1.00 |
| LCL: BS- PDS+ | 4283 | 3858 | 1.11 | 1.00 |
| LCL: BS+ PDS- | 2529 | 4530 | **0.56** | 1.50E-756 |
| Fib: BS- PDS- | 14657 | 6726 | **2.18** | 1.34E-7176 |
| Fib: BS+ PDS+ | 8278 | 5799 | **1.43** | 1.36E-1154 |
| Fib: BS- PDS+ | 3491 | 5304 | **0.66** | 7.30E-858 |
| Fib: BS+ PDS- | 2418 | 6152 | **0.39** | 1.47E-1765 |

BS+ and BS−: regions with significantly increased or decreased chromatin accessibility in BS compared to WT; PDS+ and PDS−: regions with significantly increased or decreased chromatin accessibility in PDS compared to control. Fold enrichment values in bold denote *P* < 0.05 of the corresponding hypergeometric tests.

**Supplementary Table 3. Hypergeometric tests of enrichment and depletion of overlapping differentially expressed genes upon BLM deficiency and PDS treatment.**

| DE gene catogories | Observed | Expected by chance | Fold enrichment | *P* (Hypergeometric test) |
| --- | --- | --- | --- | --- |
| LCL: BS- PDS- | 770 | 622 | **1.24** | 2.15E-13 |
| LCL: BS+ PDS+ | 721 | 646 | **1.12** | 1.23E-131 |
| LCL: BS- PDS+ | 467 | 548 | **0.85** | 3.25E-05 |
| LCL: BS+ PDS- | 256 | 733 | **0.35** | 1.35E-04 |
| Fib: BS- PDS- | 1408 | 859 | **1.64** | 3.89E-133 |
| Fib: BS+ PDS+ | 872 | 760 | **1.15** | 6.69E-66 |
| Fib: BS- PDS+ | 632 | 824 | **0.77** | 4.02E-20 |
| Fib: BS+ PDS- | 443 | 793 | **0.56** | 7.78E-08 |

BS+ and BS−: genes with significantly increased or decreased expression in BS compared to WT; PDS+ and PDS−: genes with significantly increased or decreased expression in PDS compared to control. Fold enrichment values in bold denote *P* < 0.05 of the corresponding hypergeometric tests.

**Supplementary Table 4. Mutations in *BLM* gene and medical records of the Bloom Syndrome family members.**

| Sample ID | Sex | Relation to proband | Bloom syndrome status | *BLM alleles* | Zygosity | Age at donation (month) | Neoplasm |
| --- | --- | --- | --- | --- | --- | --- | --- |
| FBL131 | M | Father | Carrier | *blm*^Ash^/+ | Heterozygous | 456 |  |
| FBL132 | F | Mother | Carrier | *blm*^Ash^/+ | Heterozygous | 446 |  |
| FBL305 | M | Patient 1 | Affected | *blm*^Ash^/ *blm*^Ash^ | Homozygous recessive | 19 | Wilm's tumor at 36 months, AML at 72 months, myelodyplastic syndrome at 72 months |
| FBL306 | F | Patient 2 | Affected | *blm*^Ash^/ *blm*^Ash^ | Homozygous recessive | 63 | None |
| FBL301 | M | Brother | Wild-type | +/+ | Homozygous | 171 |  |
| FBL302 | F | Sister | Carrier | *blm*^Ash^/+ | Heterozygous | 151 |  |

*blm*^Ash^*:* c.2207_2212delATCTGAinsTAGATTC in exon10 in BLM.

**Supplementary Table 5. Primer sequences used in G4 ChIP qPCR.**

| Primer | Sequence | G4-ChIP peak | G4 motifs |
| --- | --- | --- | --- |
| KIF14_G4_for | CGGTAGCCGTCTCTGAATG | + | + |
| KIF14_G4_rev | CTTTAGCAGAACCCGAGGAG |  |  |
| RPA3_G4_for | CGGAAGTTGACAGATACAGGG | + | + |
| RPA3_G4_rev | GATCGCAGAAAGGTAGTCTCAG |  |  |
| GAPDH_G4_for | GCTACTAGCGGTTTTACGGGCG | + | + |
| GAPDH_G4_rev | TGCGGCTGACTGTCGAACAGG |  |  |
| HTR6_G4_for | GGCGATTTGTCCAATATTTCCC | - | + |
| HTR6_G4_rev | CTGTGACCTGCCCTTATCC |  |  |
| ESR1_for | GAAACAGCCCCAAATCTCAA | - | - |
| ESR1_rev | TTGTAGCCAGCAAGCAAATG |  |  |
| TMCC1_for | GTGGTACACTGCCTACAGTATT | - | - |
| TMCC1_rev | GTATAACGCCTGGGCTATGT |  |  |
